# Flux–species sampling enables holistic exploration of biocircuit behavior

**DOI:** 10.64898/2026.07.21.739804

**Authors:** Qinguo Liu, Wenqin Zhou, Xinying Ren, John Paul Marken, Fangzhou Xiao

## Abstract

Biocircuit behavior is usually discovered by drawing uncertain kinetic parameters and simulating, but the scored phenotypes are functions of steady-state species, catalytic fluxes, dominance relations, and local response matrices. We introduce flux–species sampling (FSS), which samples that functional layer directly and algebraically inverts to parameters. This coordinate change turns regulatory modes into simple dominance chambers, gives every full-dimensional behavior-realizing regime positive measure, and can replace the sampling stage of existing pipelines without changing downstream stability or phenotype tests. A sequestration toy shows how a large species regime can collapse into a narrow parameter strip. In an enzymatic feedback motif, FSS rescues free-*A*\* adaptation missed by parameter sampling and identifies the regime as non-Michaelian and complex-dominated. In MultiFate-*N*, FSS separates a sampling limitation from a structural limitation and rescues parameter sets realizing all 2^*N*^ states.

**Highlights:**

- Flux–species sampling (FSS) draws the control/function layer where biocircuit behavior is read
- FSS boxes intersect every full-dimensional behavior-realizing regime with positive measure
- FSS is no slower than parameter-space sampling and is a hot fix for existing pipelines
- FSS rescues non-Michaelian free-*A*\* adaptation missed by parameter sampling
- FSS rescues the all-2^*N*^ symmetric MultiFate regime missed by fixed parameter boxes

**eTOC blurb:** This study reframes biocircuit exploration by sampling steady-state species and fluxes rather than raw parameters. The resulting flux–species coordinate exposes regulatory dominance regimes, rescues adaptation and multistability regimes missed by parameter sampling, and gives finite sampling nulls an explicit scope.

## 1 Introduction

Mechanistic models are central to systems and synthetic biology because they connect circuit architecture to behavior ^1^. They are used not only to fit data, but also to explain mechanisms, test design principles, and guide experiments. A recurring question is therefore architectural: what behaviors can a given biomolecular circuit realize? Answering this question is difficult because kinetic parameters are often uncertain over many orders of magnitude, nonlinear steady-state equations can have multiple solution branches, and closed-form analysis is rarely available. As a result, numerical parameter sampling has become a default exploratory strategy: choose a parameter box, draw parameter sets, solve or simulate the model, and score the resulting behavior.

A successful parameter sample is immediately informative. It demonstrates that the circuit can realize the behavior, and the corresponding operating point can be dissected to identify a mechanism. A failed search is much harder to interpret. If no sample realizes the phenotype, the behavior may indeed be structurally impossible for the circuit. But the realizing region may also lie outside the chosen parameter box, or occupy a small, thin, or ill-conditioned subset within it. In a 12-dimensional log-parameter cube, even 10^6^ samples correspond to only 10^6*/*12^ ≈ 3 samples per axis. Thus, a null result from a fixed parameter box is not, by itself, evidence of architectural impossibility.

The problem is not only dimensionality. It is also a problem of coordinates. Mechanistic models are written in kinetic parameters, whereas the behaviors we score are read after solving the model: from steady-state free and bound species, catalytic fluxes, local response coefficients, and stability eigenvalues. Parameter sampling therefore generates candidates in one coordinate system and evaluates them in another. The same dominance regime that is simple in steady-state flux–species coordinates can become a compound set of parameter inequalities, involving products such as *K*_1_*K*_2_ ≫ *q*_1_*K*_4_ or near-equalities among totals. Consequently, a failure of parameter-space search can be mistaken for a mechanistic limitation of the circuit.

This mismatch is mechanistic, not only geometric. The steady-state behaviors we interpret are organized by dominance regimes: which molecular forms are free or bound, which catalytic fluxes balance or dominate, and which terms control the local response. These regimes are first-order comparisons among steady-state species and fluxes. In log coordinates, conditions such as *C* ≫ *A, v*_1_≫ *v*_2_, or *αA* ≫ *βB* are displayed partial orders. Thus the natural coordinate for the object we want to discover is not the raw parameter box, but the dominance chamber in flux–species space.

Here we develop flux–species sampling (FSS), a complementary strategy that reverses the usual direction of search. Instead of drawing kinetic parameters and asking which steady state and dominance regime they produce, FSS draws steady-state molecular species and admissible catalytic fluxes, algebraically inverts these quantities to compatible kinetic parameters, and then applies the same positivity, stability, and phenotype tests used in conventional parameter sampling. We prove that FSS is holistic in a precise regime-level sense: with a box width set by the desired dominance margin, its flux–species box intersects every full-dimensional dominance regime of the network. This coordinate change lets us rescue missing behaviors behind negative conclusions from parameter-space sampling. In the enzymatic feedback motif of Ma et al. ^8^ and Jeynes-Smith and Araujo ^4^, FSS rescues free-*A*\* adaptation, but in a non-Michaelian, complex-dominated regime whose setpoint is shifted away from the Michaelian value. In the MultiFate-*N* network of Zhu et al. ^13^, FSS rescues parameter sets realizing all 2^*N*^ symmetric fate states across the tested sizes, showing that the collapse seen in parameter-space searches is a sampled-region limitation rather than a structural capacity limit.

We first use a single binding reaction to introduce the coordinate change and show the simplest distortion: a clear species-dominance wedge becomes a thin, ill-conditioned strip in parameter coordinates. The two case studies then show the constructive consequence: FSS does not merely say that a parameter-space miss is inconclusive, but recovers the missed behavior and the dominance conditions that make it possible.

## 2 Results

### 2.1 A toy network exposes the coordinate problem

Before introducing the general framework, we start with the smallest network in which the coordinate issue is visible by algebra alone. The toy is a single binding reaction, *A* + *B* ⇌ *C*, in which free *A* gates a downstream readout flux *v* = *k*_cat_*A* (Fig. 1a,b). The same motif appears in ligand– receptor binding ^5^, transcription-factor titration and protein sequestration^2,3^, and miRNA/small-RNA sequestration^6,9^. This network can be parameterized either by totals and binding constant, (*A*_tot_, *B*_tot_, *K*), or by the species themselves, (*A, B, C*), with

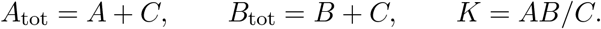

**Figure 1.**
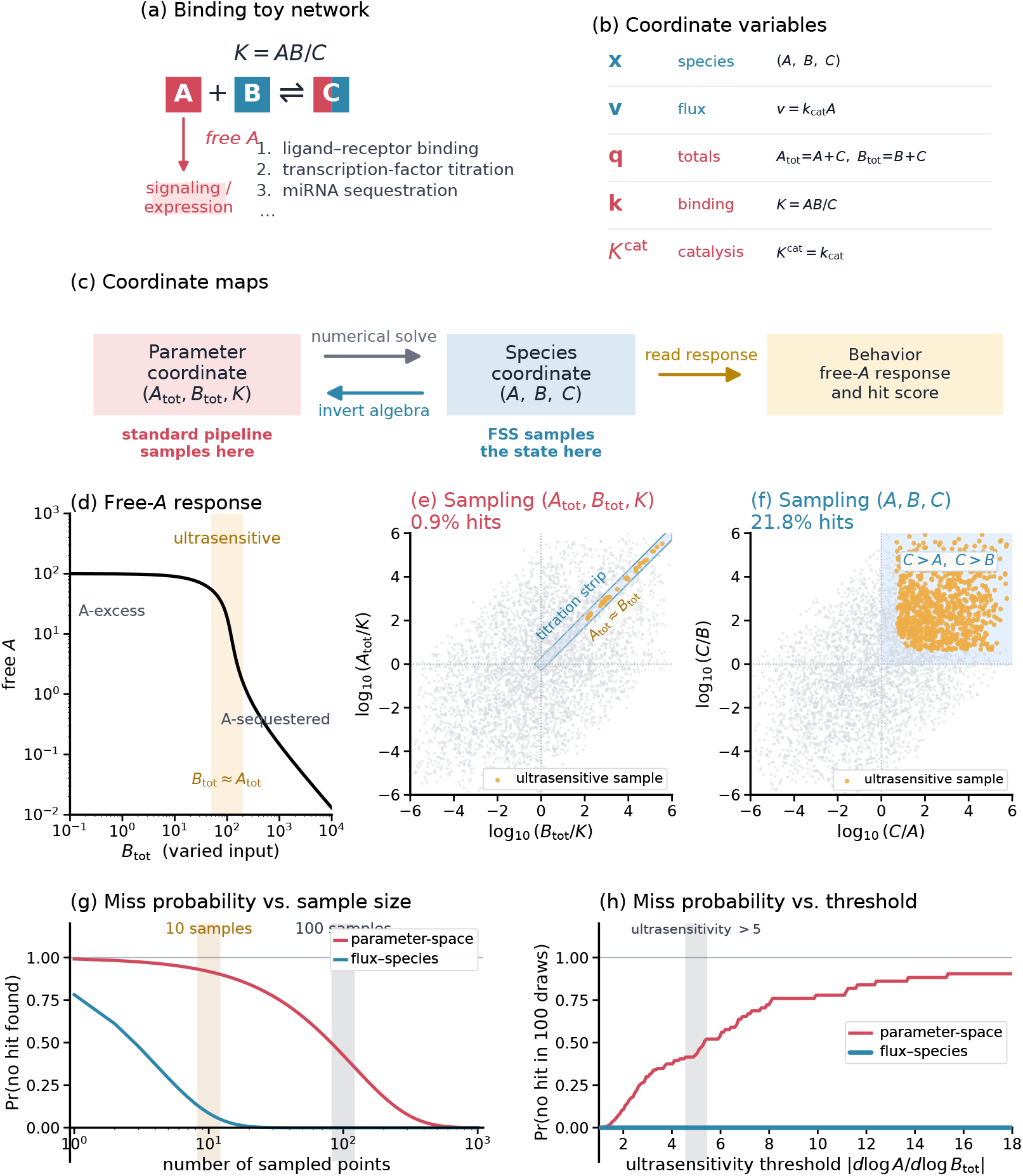
Coordinate dependence in a three-species toy. (a) A binding reaction *A* + *B* ⇌ *C* gates a downstream readout through free *A*. (b) The same state can be described by species ***x*** = (*A, B, C*), totals (*A*_tot_, *B*_tot_) = (*A* + *C, B* + *C*), and binding constant *K* = *AB/C*. (c) Parameter sampling first solves for a species state; species-coordinate sampling draws that state directly and inverts algebraically to compatible parameters. (d) Free *A*, and therefore *v* = *k*_cat_*A*, changes sharply as *B*_tot_ crosses *A*_tot_. (e) Parameter-space sampling of (*A*_tot_, *B*_tot_, *K*), shown in (*A*_tot_*/K, B*_tot_*/K*). Points passing the ultrasensitivity criterion *d* log *v/d* log *B*_tot_ *>* 5 lie near *A*_tot_ ≈ *B*_tot_ ≫ *K*. (f) Species-coordinate sampling of (*A, B, C*), shown in *C/A, C/B*). The same criterion selects the *C*-dominant region. (g) Probability that *m* draws return no hit at the fixed threshold *>* 5. (h) Probability that 100 draws return no hit as the ultrasensitivity threshold is varied.

We ask for a switch-like response of the readout to the titrating input *B*_tot_ (Fig. 1d). Since *k*_cat_ is fixed in this toy, this is equivalently a large local derivative of free *A*:

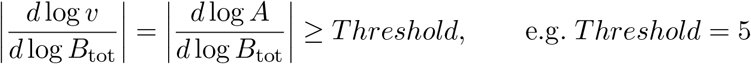

We then sample the same behavior in the two coordinate systems. In parameter coordinates, sampling (*A*_tot_, *B*_tot_, *K*) finds ultrasensitive points only in a thin strip near *A*_tot_ ≈ *B*_tot_ ≫ *K*, with a per-draw hit rate of about 1% (Fig. 1e). In species coordinates, sampling (*A, B, C*) finds the same behavior throughout the *C*-dominant region, *C* ≫ *A* and *C* ≫ *B*, with a per-draw hit rate of about 23% (Fig. 1f). The circuit and phenotype test are unchanged; only the coordinate used to generate candidate states has changed. The coordinate determines the probability that finite sampling encounters the behavior. We estimate the per-draw hit probability *p* in the two coordinates (Fig. 1e,f), and then plot the corresponding probability that *m* finite draws miss the behavior, (1 − *p* )^*m*^ (Fig. 1g,h). Thus, a thin parameter-space image can turn an existing behavior into a high-probability sampling miss, which can then be mistaken for a structural negative.

This toy exposes the coordinate problem: a real behavior can occupy a thin or ill-conditioned image in parameter space. The next subsection introduces the general flux–species framework and the sampling recipe that searches directly in the coordinates where such dominance regimes are read.

### 2.2 Flux–species sampling: framework and recipe

We decompose a biomolecular circuit into two kinds of reactions: *binding* reactions (fast, reversible, mass-action) and *catalysis* reactions (slow, irreversible production or degradation). Following Xiao ^10^, Xiao et al. ^12^ we collect them in a differential-algebraic system,

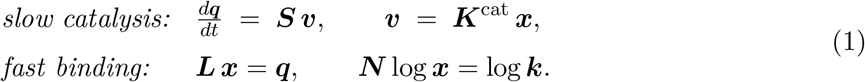

where 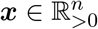 is the *species-concentration* vector, 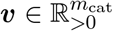 the *catalytic fluxes* vector, and 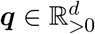 the totals (conserved quantities of the binding layer). The matrices ***L* ∈ ℝ** ^*d×n*^ (conservation laws), ***N* ∈ ℝ** ^*r×n*^ with *r* = *n − d* (reduced binding stoichiometry), 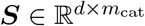 (catalysis stoichiometry), and 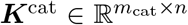 (catalysis-rate matrix; entry 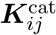 is the rate constant for the *i*-th catalysis reaction having *x*_*j*_ as an active species) are combinatorial properties of the network topology.

The fast-binding block contains mass conservation and binding equilibrium in log form (mass-action 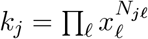), and together they uniquely fix the positive species vector ***x*** for the mass-action binding networks considered here. The slow-catalysis block says that the catalytic fluxes ***v*** = ***K***^cat^ ***x*** drive the slow evolution of ***q***.

The toy model above is the binding-only instance of this structure; the Supplemental Information formulates the toy and the two case-study networks in the same flux–species language in §S1.3. *The flux–species pair* (***v, x***) *is the coordinate FSS samples*: it is the immediate layer from which behavior is read, with the parameters (***q, k, K***^cat^) recovered from it by inversion (§S3.2), generalising the species-coordinate draw shown in the binding toy (Fig. 1c).

#### Dominance regimes are the behavioral coordinates

Many reduced steady-state descriptions are leading-order descriptions of dominance regimes: they follow after specifying which species dominate conserved pools and which fluxes dominate production or degradation terms. The toy already shows this. The binding network *A* + *B* ⇌ *C* has four regimes, determined by whether *A* or *C* dominates *A*_tot_ = *A* + *C*, and whether *B* or *C* dominates *B*_tot_ = *B* + *C* (Fig. S1; see §S2). The switch-like response in Fig. 1d requires the *C*-dominant regime, *C* ≫ *A* and *C* ≫ *B*; we show in §S6.3 how this dominance gives a large local gain |*d* log *A/d* log *B*_tot_|.

The enzymatic module *E* + *S* ⇌ *C* → *E* + *P* illustrates the same idea. Its product-forming flux is *v* = *k*_cat_*C*, while fast binding and conservation give *E S* = *KC, q*_*S*_ = *S* + *C*, and *q*_*E*_ = *E* + *C*. Once a dominance regime is specified, the leading-order behavior follows. In the free-substrate regime *S* ≫ *C, q*_*S*_ ≈ *S*, giving the usual Michaelis–Menten reduction. In the complex-dominated regime *C* ≫ *S*, the same conservation law gives *q*_*S*_ ≈ *C*, so the product flux scales as *v* ≈ *k*_cat_*q*_*S*_.

Generalising, the *bioregulatory mode* of a binding network at any state is determined by its *dominance profile*—the matrix whose (*i, j*)-th entry is the fractional contribution of species *j* to total *i*,

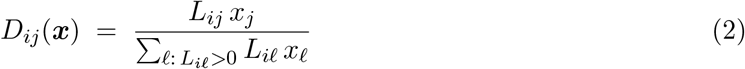

as developed in §S2 and §S2^10–12^. Reaction orders, ultrasensitivities, regulatory motifs, and the local linearisation of the dynamics all read off this dominance structure (§S5). When *D*_*ij*_ ≈ 1 for one species *j* in each total *i*, the network admits a small-form reduced description fixed by a few asymptotic inequalities in log ***x***.

With catalysis, the same idea applies to the production and degradation terms selected by ***S***: which flux dominates each birth or death side of each moving total is the catalytic dominance layer. The binding and catalytic dominance layers together form the bioregulatory mode, and both are read directly from (***v, x***). Parameters determine which flux–species state is realised, but once (***v, x***) is fixed, the mode is determined by comparisons among species and fluxes themselves. The full two-layer dominance construction is given in §S2, and the flux–species coordinate chart is given in §S3.

This is the coordinate used by the sampler: draw the operating point where regimes and phenotypes are read, then invert to compatible parameters and apply the same downstream tests.

#### The sampling recipe

Given a network in form (1), FSS proceeds as follows (Fig. 2):

1. Sample the flux–species state (***v, x***) ∼ Uniform(log -box) (for a binding-only circuit, just ***x***).
2. Compute ***k*** = exp(***N*** log ***x***) (*r* scalar evaluations).
3. Compute ***q*** = ***Lx***. Split ***q*** = (***q***^cat^, ***w***) where ***w*** are conserved on *both* binding and catalysis time-scales (system inputs, totally conserved totals) and ***q***^cat^ evolves under catalysis.
4. Recover ***K***^cat^ from ***v*** = ***K***^cat^***x***: column-specificity (one flux per species) makes this explicit, and the independent fluxes—the free coefficients left by the balance ***SK***^cat^***x*** = 0—are precisely the ***v*** half of the draw, sampled log-uniformly within the nonzero pattern.
5. Compute the analytic Jacobian (see §2.2) at the sampled ***x***.
6. Read off any dynamical property of interest at this fixed point — stability, adaptation precision, ultrasensitivity, robustness — as a closed-form algebraic function of ***x*** and the Jacobian.
7. (Optional) Reject samples whose implied (***k, K***^cat^, ***q***) violate biological or experimental priors.

**Figure 2.**
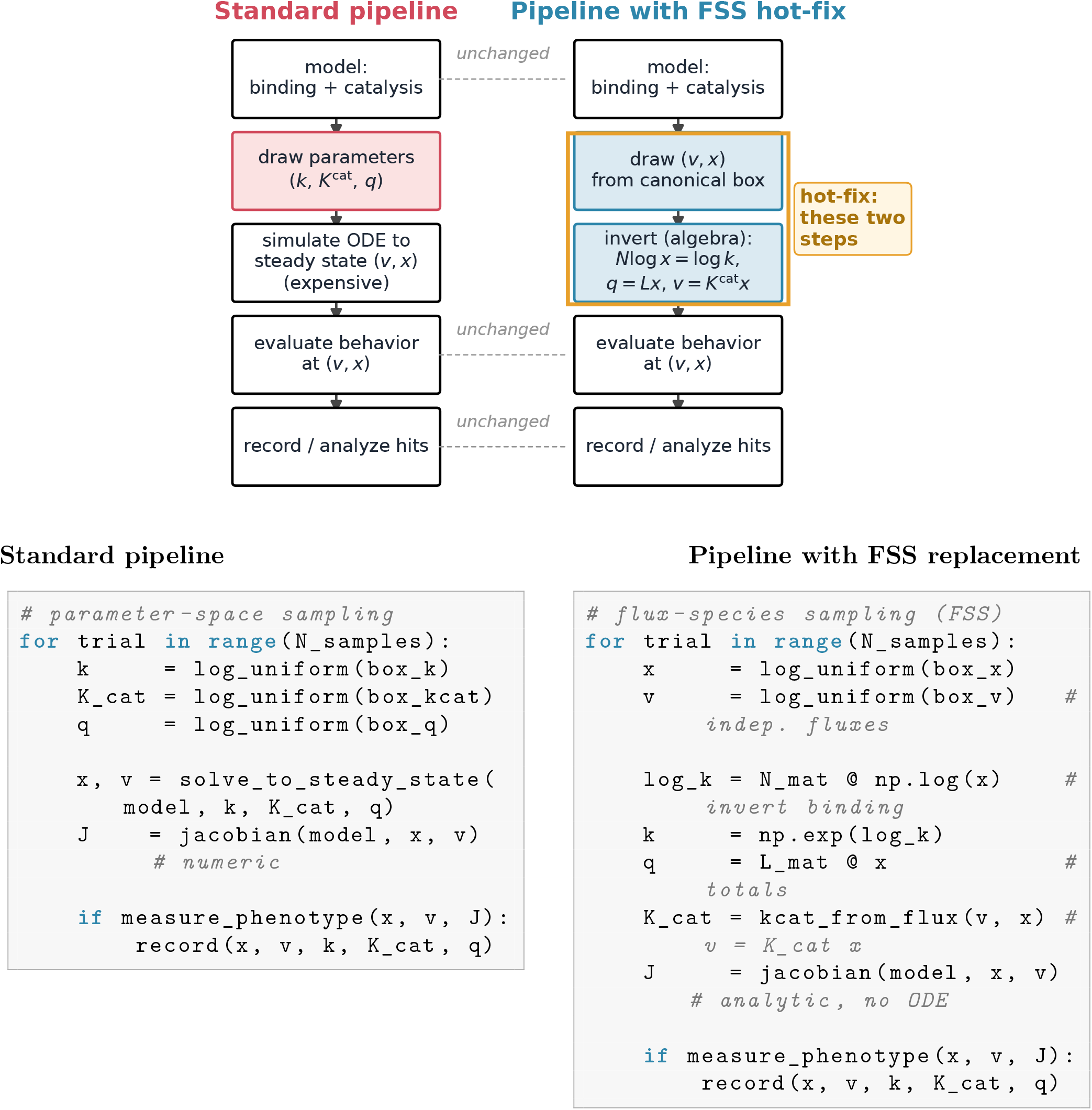
FSS replaces one stage of the standard pipeline. *Top:* the standard pipeline draws parameters from a chosen box and integrates an ODE to steady state to find ***x***. FSS draws the flux–species state (***v, x***) directly from a canonical log-box and *inverts* the binding and catalysis equations to recover the parameters (the schematic shows the binding-only case, where ***v*** is absent and ***x*** alone is drawn); model specification, behavior measurement, and hit recording are unchanged. *Bottom:* the corresponding pseudo-code diff. A few lines of linear algebra replace the sample-and-simulate block and provide the analytic Jacobian used for local scoring.

Steps 1–6 are purely algebraic; no ODE integration is required. Figure 2 shows both the pipeline replacement and the minimal code-level change: the standard draw-then-simulate stage is replaced by draw-(***v, x***)-then-invert, while the phenotype measurement and downstream hit recording stay unchanged.

#### Stability, sensitivity, and other phenotypes are algebraic in *x*

Differentiating the binding-equilibrium and conservation constraints with ***k*** held fixed gives a single linear system whose inverse encodes the response of every species to any total or input perturbation. From it the Jacobian of the slow dynamics on ***q***^cat^ at any sampled ***x*** follows in closed form:

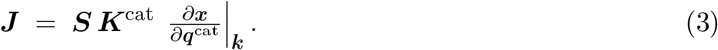

Stability is a *d*_cat_ *× d*_cat_ eigenvalue test of ***J*** ; the input–output transfer function uses the same matrix to deliver adaptation precision, sensitivity, and ultrasensitivity-type phenotypes via the implicit function theorem applied to the catalysis flux balance. Full derivations are in Supplemental Information, §S5. Thus *every dynamical property considered here at a fixed point is a closed-form algebraic function of the sampled* ***x***: the “simulate–perturb–repeat” loop of the standard pipeline collapses to a single matrix inversion.

#### FSS is a drop-in replacement: only one pipeline stage changes

The two case studies below use standalone FSS implementations, but this overstates the adoption cost. Every biocircuit modelling script that does numerical sampling already contains the ingredients FSS needs: the binding network stoichiometry ***N***, the conservation matrix ***L***, and the catalysis stoichiometry ***S*** are combinatorial properties of the species topology that any mechanistic model encodes (explicitly or implicitly). Replacing the sample-and-simulate loop with FSS touches one stage of the pipeline and leaves everything else untouched (Fig. 2, bottom). In our reference implementation, the substitution was ∼ 30 lines of Python on each case study.

#### FSS does not add per-sample computational cost

Per sample, FSS does an *O*(*n*^2^) linear solve (the binding response matrix) plus a few *O*(*n*) matrix–vector products (steps 2–4). The standard pipeline does an ODE solve to steady state, which is at best *O*(*n*^2^) per step and typically many steps, often with stiffness from the disparate binding and catalysis time-scales. Empirically, on the adaptation and MultiFate case studies below, the per-sample wall-clock cost of FSS is comparable to or smaller than the standard pipeline. FSS therefore does not trade sampling scope for higher per-sample runtime.

### 2.3 Completeness behind flux–species sampling

Dominance regimes are the behavioral coordinates on which completeness rests. Within a dominance regime, the leading species carriers, leading flux balances, and local response structure are fixed, so the reduced behavior is constant to leading order throughout the chamber. The regime itself is specified by first-order comparisons between like quantities—species against species and flux against flux. In log flux–species coordinates, each comparison is a hyperplane with a stoichiometry-set offset, and the regimes are chambers of the resulting difference arrangement. Completeness therefore means that a canonical flux–species box intersects every full-dimensional behavioral chamber and gives positive measure once the box is wide enough for the requested dominance margin.2

#### Theorem 1

(Canonical completeness). *Let β* = max_*i*_ max_*j,ℓ*_ log(*L*_*iℓ*_*/L*_*ij*_) *be the network’s stoi-chiometric spread and* 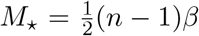 . *For every M > M*_⋆_, *the species box* ( − *M, M* )^*n*^ *in* log ***x*** *meets the interior of* every *dominance regime. To additionally realise a quantitative strength R (a precision, fold change, or margin), it suffices that M > M*_⋆_ + log *R*.

Theorem 1 is a coordinate-level statement about dominance chambers in species space (§S4). Parameter coordinates generally represent the same chambers by compound inequalities and near-equality constraints among heterogeneous parameters, rather than by a topology-set canonical box.

With catalysis, the same guarantee holds in the full flux–species coordinate (***v, x***); the radius is then controlled by the stoichiometry of both the binding and birth–death layers (§S4.2, Theorem S1).

#### Theorem 2

(Finite-sample misses). *Draw m species points log-uniformly from a box satisfying Theorem 1. If a regime-determined behavior is realisable with per-draw probability p >* 0 *in that box, the probability that all m draws miss it is* (1 − *p*)^*m*^.

Theorem 2 makes a finite-sampling null explicit in the coordinate where the behavior-realizing regimes are sampled. Once the flux–species box, dominance margin, and phenotype criterion have been declared, an empty FSS run means that no draw in that declared box met the criterion within the finite budget. If the target behavior occupies a positive-measure regime with per-draw probability *p*, the miss probability is (1 − *p*)^*m*^; for displayed partial orders, the chain-length certificate in §S4.3 gives a conservative lower bound *p*_min_ for the rarest displayed chamber, hence an upper bound (1− *p*_min_)^*m*^ on the miss probability.

The central consequence is constructive. FSS is not used merely to argue that a parameter-space miss is not a proof of impossibility; it is used to rescue misses caused by coordinate distortion. When a behavior is realised on a full-dimensional dominance chamber that becomes thin, shifted, or ill-conditioned in a raw parameter box, FSS samples that chamber directly, returns hits, and reports the dominance profile of the recovered mechanism. Biological or experimental plausibility can then be imposed afterward by filtering the recovered parameters (§S7).

Together, the construction and the two guarantees make FSS complete and holistic at the level needed for exploratory mechanism search: it samples the coordinates in which behavioral regimes are chambers, records finite-sample miss probabilities in that declared box, and returns the dominance profile of each hit for mechanism interpretation. We next use two case studies where this coordinate change rescues behaviors that parameter-space sampling had made look absent or marginal.

### 2.4 Case study 1: competitive binding does not preclude adaptation

#### 2.4.1 The controversy: Michaelian adaptation versus explicit complexes

Ma et al. ^8^ asked which three-node enzymatic circuits *adapt*: after a step input, the output should transiently respond and then return to its pre-stimulus level. Their screen quantified this by an adaptation precision |*d* log *O/d* log input | ^−1^ *>* 10 together with a non-trivial transient sensitivity. The crucial modelling assumption was Michaelis–Menten kinetics: enzyme–substrate complexes were assumed negligible compared with the free substrate species retained in the reduced equations. Under that assumption, Ma et al. identified the negative-feedback architecture in Fig. 3a: input *I* activates *A, A* activates *B, B* feeds back to inactivate *A*, and a constitutive enzyme *E* deactivates *B*\*. In the Michaelian reduction, the free active output *A*\* tracks the input-independent setpoint

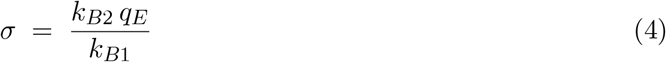

where activated forms are denoted by *.

**Figure 3.**
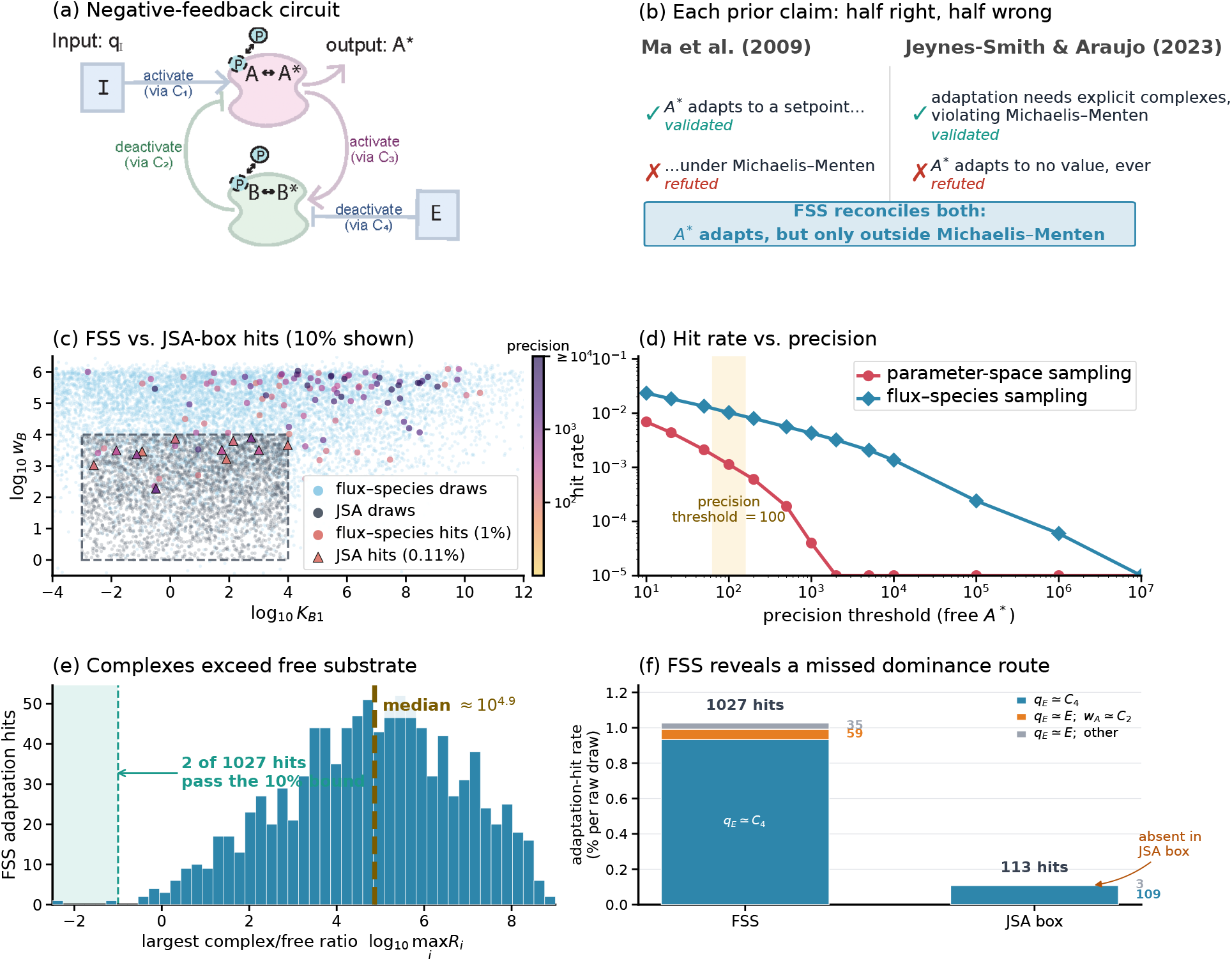
FSS recovers the free-*A*\* adaptation that Jeynes-Smith & Araujo declared impossible “under any parametric conditions”. (a) Negative-feedback topology with explicit complexes *C*_1_− *C*_4_. (b) The two prior claims, scored. Ma et al. ^8^ : free *A*\* adapts to *some* setpoint (validated) but only under a Michaelian reduction (refuted—the adapting regime is complex-dominated). Jeynes-Smith and Araujo ^4^ : adaptation requires non-negligible complexes (validated) but free *A*\* adapts to *no* setpoint, ever (refuted). FSS reconciles both: free *A*\* adapts in complex-dominated regimes, including a *C*_4_-route expression *A*\* *σK*_*B*1_*/B* and the free-*B*-buffered subcase *A*\* ≈ *σK*_*B*1_*/w*_*B*_. (c) Adaptation hits (precision ≥ 100, responsive, stable) in (log_10_ *K*_*B*1_, log_10_ *w*_*B*_): FSS realises them at 1.0% (circles), the JSA box at 0.11% (triangles); faint points show one tenth of each 10^5^ -draw cloud, hit colour encodes 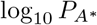. (d) Hit rate versus precision threshold: the FSS–box gap widens as the demand tightens. (e) The adaptation lives where the Michaelian reduction *fails*. For each FSS adaptation hit we plot the largest log complex/free ratio among {*C*_1_*/A*, (*C*_2_ + *C*_3_)*/A*\*, *C*_3_*/B*, (*C*_2_ + *C*_4_)*/B*\*} The negligible-complex condition would place all hits left of 1, meaning every explicit complex pool is below 10% of its corresponding free substrate; almost all hits violate that bound. (f) Dominance-route split of adaptation hits into *q*_*E*_ ≈ *C*_4_, *q*_*E*_ ≈ *E*; *w*_*A*_ ≈ *C*_2_, and *q*_*E*_ ≈ *E* other. The free-*E, C*_2_-buffered route appears in FSS but is absent from reconstructed JSA-box hits. Direct Michaelian-complex-estimate error and fixed/route setpoint-error diagnostics are shown in Figs. S7 and S9.

Jeynes-Smith and Araujo ^4^ challenged this conclusion by making the protein–protein complexes explicit. The competitive-binding model contains the four binding equilibria

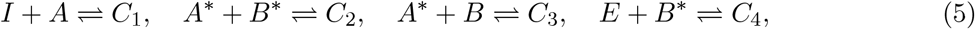

with dissociation constants *K*_*A*1_, *K*_*A*2_, *K*_*B*1_, *K*_*B*2_ and slow catalysis dynamics

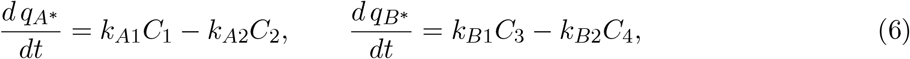

where 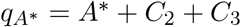 and 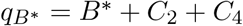 are the active-form totals; *w*_*A*_, *w*_*B*_, *q*_*I*_, *q*_*E*_ are conserved moieties (Supplemental Information, §S8.3). Their mechanistic objection was correct: explicit complexes are generally not negligible in this network, as the FSS hits later show directly in Fig. 3e. Their sampling conclusion went further. From 3 × 10^5^ log-uniform draws over the parameter box

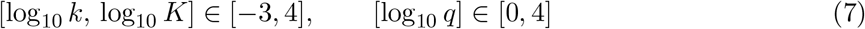

(the Jeynes–Smith–Araujo, or JSA, range), they found no adapting free-*A*\* set and interpreted this as a universal negative: free *A*\* would not track *σ*, or any other non-trivial setpoint, for any biochemical rate constants or protein abundances. Figure 3b summarizes the resulting controversy: Ma et al. were right that the architecture can adapt, but their Michaelian assumption misses the actual regime; Jeynes-Smith and Araujo were right that explicit complexes invalidate the Michaelian reduction, but their parameter-space search missed free-*A*\* adaptation itself.

#### 2.4.2 Resolving the controversy: flux–species sampling finds the adaptation

##### FSS revises the parameter-sampling negative

Scoring free-*A*\* adaptation by precision | *d* log *A*\**/d* log *q*_*I*_| ^−1^ ≥ 100, responsiveness (Supplemental Information, §S5.7), and stability, all evaluated from the sampled state via the Jacobian (3)—FSS finds free-*A*\* adaptation in 1027/10^5^ = 1.0% of draws, compared with 113/10^5^ = 0.11% for the JSA parameter box (Fig. 3c). The gap widens as the demanded precision increases (Fig. 3d), making the finite-sample miss much less likely under FSS.

Diving into the FSS hits shows why the result is not Michaelis–Menten adaptation in disguise. The Michaelian reduction assumes enzyme–substrate complexes are negligible compared with their free substrate pools. For each FSS adaptation hit, we compare *C*_1_, *C*_2_ + *C*_3_, *C*_3_, and *C*_2_ + *C*_4_ with *A, A*\*, *B*, and *B*\*, respectively. Only 2 of the 1027 hits satisfy the 10% negligible-complex bound; the median largest complex/free ratio is 10^4.87^ (Fig. 3e). Thus Ma et al. were right about adaptation but wrong about the Michaelian regime, whereas Jeynes-Smith and Araujo were right that the Michaelian assumption fails but wrong that free *A*\* cannot adapt (Fig. 3b).

##### Dominance routes reveal the missed mechanism

The hit-rate comparison is only the first layer of the result. Because each FSS hit is a complete steady-state species vector, the hit set can be read as a collection of dominance regimes. This is where the missing mechanism becomes visible: a regime that is an ordinary species-composition class can map to a thin, coordinated strip in raw parameter coordinates.

Figure 3f splits the adaptation hits into three routes. The common route has *q*_*E*_ ≈ *C*_4_: it accounts for 933*/*1027 FSS hits and 109 of the 113 JSA-box hits. The exact *B*-arm balance gives

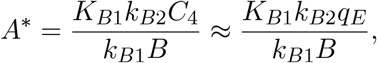

so adaptation follows when the effective free-*B* denominator is buffered against *q*_*I*_. The simple subcase *B* ≈ *w*_*B*_, analyzed explicitly by Liu et al. ^7^ and summarized in §S8.4, gives

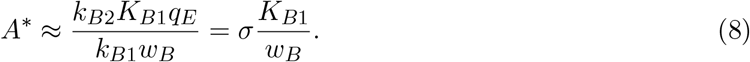

The setpoint is therefore input-independent even though it is generated by an explicit-complex, non-Michaelian regime.

The route missed by the parameter box has *q*_*E*_ ≈ *E* and *w*_*A*_ ≈ *C*_2_: it contains 59 FSS hits but 0 reconstructed JSA-box hits. In this route *C*_4_ = *EB*\**/K*_*B*2_ ≈ *q*_*E*_*B*\**/K*_*B*2_, so the buffered object is *B*\**/B*, or equivalently *C*_2_*/B* when *C*_2_ dominates the active pools:

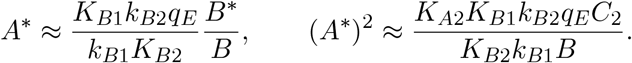

If *C*_2_ ≈ *w*_*A*_ and *B* ≈ *w*_*B*_, the right-hand side has no input *q*_*I*_. In species coordinates, *q*_*E*_ ≈ *E* and *w*_*A*_ ≈ *C*_2_ are open dominance inequalities. In parameter coordinates, however, the same responsive regime requires the *A*-arm balance *C*_1_ = (*k*_*A*2_*/k*_*A*1_)*C*_2_ to align with the input total:

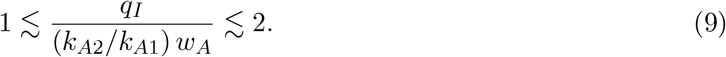

Thus the missed species-space route becomes a 0.30-decade strip in parameter space, while remaining a well-conditioned dominance chamber in flux–species space (derivation and controls in §S8.4).

Finally, the original JSA no-hit statement for free *A*\* came from a stricter setpoint-specific screen: their released code forced *A*\* = *σ*. Supplemental Information §S8.5 and Figs. S9 and S8 compare the recovered adapting hits with *σ* and dissect a fixed-*σ* JSA-box hit that also passes our response filter; together, these controls show that FSS finds real input-invariant states rather than manufacturing fixed-*σ* artifacts. That equality is a coincidence inside an adapting regime, not the generic adaptation mechanism.

### 2.5 Case study 2: MultiFate state capacity is limited by sampling, not structure

#### 2.5.1 The architecture, and what Zhu et al. saw

Zhu et al. ^13^ introduced MultiFate-*N* as an experimentally motivated, expandable mammalian cell-fate circuit. In this architecture, *N* engineered zinc-finger transcription factors share a common dimerisation domain (Fig. 4a). Each TF self-activates through its homodimer, whereas heterodimers do not bind any promoter and therefore implement mutual inhibition through the shared dimerisation substrate. The symmetric nondimensional model is

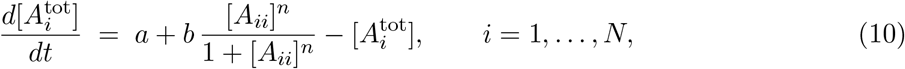

where [*A*_*i*_] is the free monomer of TF *i*, the homodimer equilibrium is [*A*_*ii*_] = [*A*_*i*_]^2^*/K*_*d*_, the heterodimer equilibria are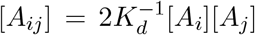, and the total concentration satisfies 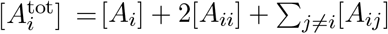. Zhu et al. ^13^ ‘s nominal “physiologically reasonable” point (their Table S1) is (*a, b, K*_*d*_, *n*) = (0.8, 20, 1, 1.5).

**Figure 4.**
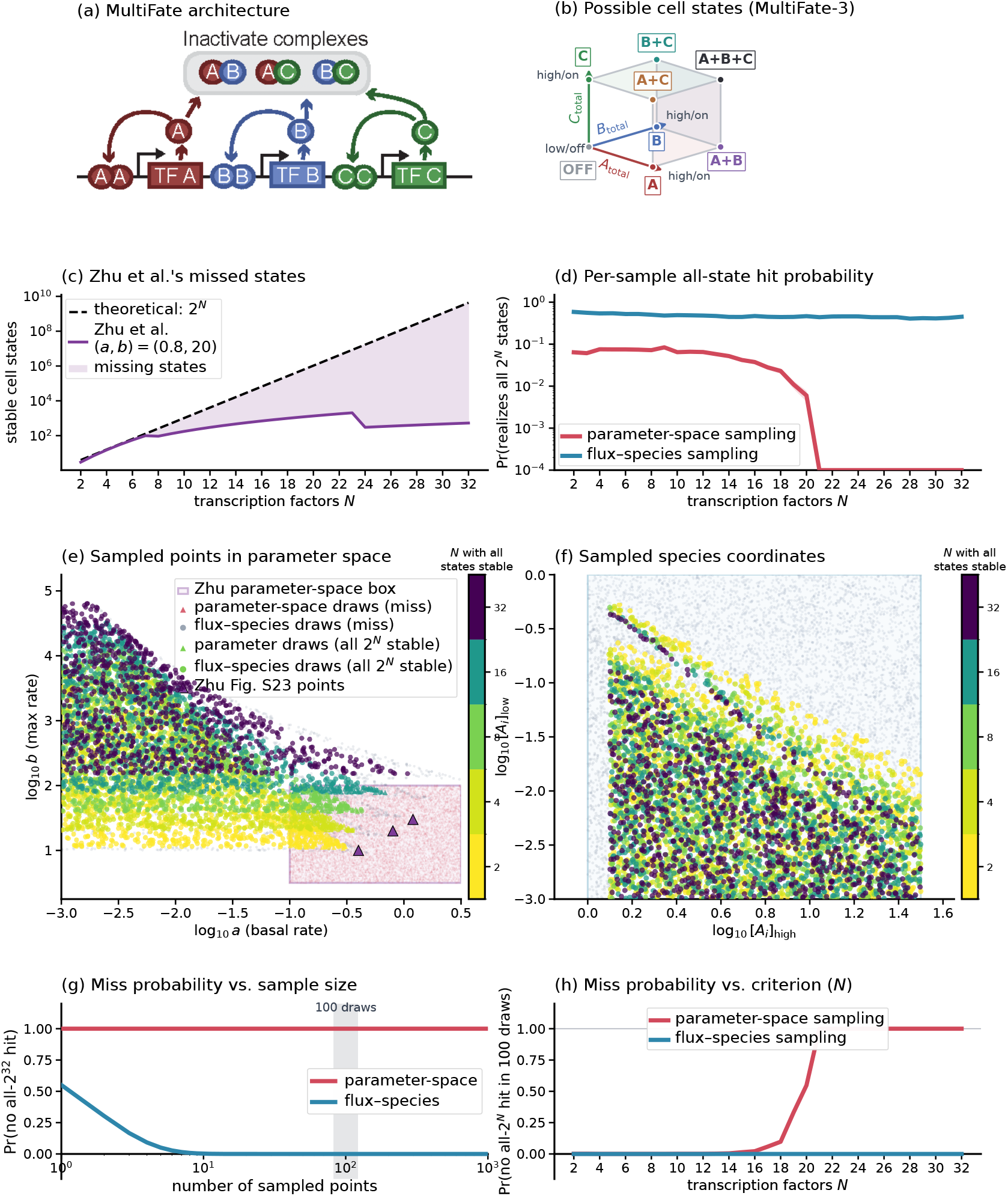
MultiFate sampling outcomes across *N* . (a) MultiFate-3 architecture. (b) The eight total-concentration cell states. (c) Zhu et al.’s nominal (*a, b*) = (0.8, 20) drops to ∼ 1.23 *×* 0^−5^% of 2^*N*^ at *N* = 32. (d) Per-sample probability of drawing a point that realizes all 2^*N*^ states, estimated from the 2000 cached draws per *N* ; lines show means and bands show binomial standard errors. (e) Parameter-coordinate plot in (*a, b*): random draws from the Zhu box are shown together with homogeneous-endpoint FSS projections. All-state FSS hits move upward in *b* as *N* increases and therefore leave the Zhu box. Colored circles and triangles are all-2^*N*^ hits, colored by *N*; gray circles and faint red triangles are misses; purple markers are Zhu Fig. S23 pairs. (f) Sampled species-coordinate plot of the same homogeneous-endpoint draws, showing that the all-state regime remains robust in the coordinates actually sampled. (g,h) Probability (1 − *p*)^*m*^ of a false negative conclusion, in the same two views as Fig. 1g,h: (g) varying the sample size *m* at a fixed criterion (all 2^32^ states, i.e. fixed *N* = 32); (h) varying the criterion (*N*, the number of fates demanded) at a fixed sample size (*m* = 100).

The self-activating homodimer module supplies the binary state variable. In the symmetric reduction, fixing the shared free-monomer pool gives a scalar steady-state equation for each TF with a low stable root and a high stable root; we call these OFF and ON, respectively (derivation in Supplemental Information, §S9.2). A MultiFate state is therefore an assignment of each TF to one of these two branches (Fig. 4b), so the architectural upper bound is 2^*N*^ stable states. Zhu et al. verified all 2^*N*^ states for *N* ≤ 3, but when they chose fixed production-rate pairs and increased *N*, the state count petered out rather than tracking 2^*N*^ (Fig. 4c). Their main-text nominal pair reached a maximum of 256 attractors at *N* = 9, and their Fig. S23 parameter pairs (*a, b*) = (0.4, 10), (0.8, 20), (1.2, 30) showed the same qualitative attrition. This makes the natural question architectural: can the MultiFate topology itself realize all 2^*N*^ states at large *N*, or did the chosen parameter region simply drift away from the all-state regime?

#### 2.5.2 Flux–species sampling finds the all-2^*N*^ regime

##### Flux–species endpoint chart

In the symmetric MultiFate model, the all-state question can be probed through a two-dimensional flux–species endpoint chart. We sample an OFF free-monomer endpoint [*A*_*i*_]_low_ and an ON endpoint [*A*_*i*_]_high_, log-uniformly on opposite sides of the Hill scale after conversion to the homodimer concentration [*A*_*ii*_] = [*A*_*i*_]^2^*/K*_*d*_. These two species endpoints impose the all-OFF and all-ON fixed-point constraints, from which the production parameters (*a, b*) are reconstructed algebraically:

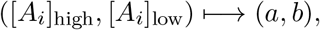

with endpoint draws that yield nonpositive parameters retained as misses. In this symmetric reduction the flux layer is implicit in the steady-state endpoint equations rather than sampled as a separate variable, but the logic is the same as in the general FSS construction: sample the steady-state species chart, reconstruct parameters, and then verify the ODE phenotype. Each reconstructed (*a, b*) is tested against all 2^*N*^ labelled states using the same Newton and Jacobian stability checks as parameter-space sampling.

The controls use the parameter coordinates directly. We evaluate Zhu et al.’s nominal (*a, b*) = (0.8, 20), and we draw 2000 log-uniform parameter samples per *N* from the Zhu box log_10_ *a* ∈ [−1, 0.5], log_10_ *b* ∈ [0.5, 2], which contains their Fig. S23 pairs. Details of the endpoint projection and verifier are in Supplemental Information, §S9.2.

##### All-state recovery

Fixed parameter choices reproduce the peter-out: Zhu et al.’s nominal pair falls to a vanishing fraction of the 2^32^ possible states (Fig. 4c), and the matched Zhu-box parameter search finds no all-state hits in 2000 draws at *N* = 32 (Fig. 4d). The failure is therefore not just a bad nominal point; it is the practical outcome of sampling a fixed parameter rectangle as *N* grows. FSS gives the opposite conclusion about the architecture. The endpoint chart rescues all 2^*N*^ states for every *N* ≤ 32 tested, and at *N* = 32 899*/*2000 raw endpoint draws (≈ 45%) are all-state hits, with invalid endpoint projections still counted as misses (Fig. 4f). Thus the same all-state criterion has a per-draw hit probability near 0.45 under FSS and zero observed hits under the matched parameter-box search (Fig. 4d; full counts in Supplemental Information, §S9.3).

##### Why fixed parameter boxes peter out

The explanation is visible after mapping the FSS hits back to (*a, b*). As *N* increases, the all-state parameter region moves upward in production strength *b*, leaving the fixed Zhu box (Fig. 4e).Therefore a fixed parameter-box search can keep sampling the same biologically motivated rectangle and still never see the all-2^*N*^ regime, even though the architecture can realise it robustly. This drift also explains Zhu et al.’s peter-out: fixed (*a, b*) choices do not track the *N* -dependent all-state region. In the flux–species coordinates actually sampled, ([*A*_*i*_]_high_, [*A*_*i*_]_low_), all-state hits remain well populated across *N* (Fig. 4f).

The same contrast gives the false-negative panels. With the criterion fixed at all 2^32^ states and the sample budget varied (Fig. 4g), FSS rapidly drives the miss probability down, while the Zhu-box empirical zero-hit estimate leaves parameter sampling at one. With 100 draws at *N* = 32, the FSS miss probability is about 10^−26^, versus one under the empirical zero-hit estimate for parameter-space sampling (Fig. 4h). The asymmetric high-dimensional stress test gives the same coordinate lesson without imposing the two-dimensional symmetric collapse (Supplemental Information, §S9.5, Fig. S11).

## 3 Discussion

### The common pattern

The three examples share one anatomy: the behavior is naturally expressed as species dominance, flux dominance, and local response in the flux–species state, while parameter sampling observes that behavior only after mapping a chosen box through the model. The main advantage of FSS is therefore not only that a negative result has a clearer scope. It is that an apparent miss can be rescued when the behavior-realizing regime is full-dimensional in flux–species coordinates but thin, shifted, or ill-conditioned in the sampled parameter box.

FSS changes the exploratory coordinate: it samples (***v, x***) where the regimes have their natural log-volume and where a hit immediately yields the dominance conditions that explain the phenotype. Because one bounded flux–species box intersects every full-dimensional dominance regime at a declared precision (Theorem 1), an FSS null has a clear scope: absent from the declared flux–species regime class and quantitative criterion, rather than merely absent from a guessed parameter box. The two case studies illustrate the stronger outcome: FSS recovers the missing adaptation route and the all-state MultiFate regime, then maps each recovered point back to the parameter conditions that made parameter-space sampling fail.

### Retrofitting and impact

Because the only required ingredients are the network matrices (***N***, ***L, S***) and a linear solver—no machine learning, symbolic algebra, or domain-specific library— FSS can be added to existing modelling code without disrupting downstream phenotype tests. In the case studies here, the replacement is localized to the sample-and-simulate block.

### Local dynamics can be captured naturally

Flux–species sampling draws (***v, x***) directly at steady state, so each accepted sample is a fixed point by construction. Local dynamic properties, including stability, responsiveness, ultrasensitivity, and adaptation precision, are then computed from the analytic Jacobian ***J*** (3), assembled from (***v, x***) with one linear solve. Stability is an eigenvalue test, and the same matrix supplies sensitivities through the implicit function theorem applied to the catalysis flux balance. Thus every fixed-point phenotype considered here is a closed-form algebraic function of the sampled state. In contrast, the standard pipeline solves an ODE to locate the fixed point and typically repeats perturbation calculations for sensitivities; FSS replaces these steps with one local matrix calculation. The scope of this paper is therefore steady-state behavior around fixed points: ultrasensitive slopes, adapted setpoints, local stability, and multistable fate states. Full transient waveforms, oscillations, and trajectory-level phenotypes are not excluded by the philosophy of FSS, but they would require sampling paths or time-indexed states rather than the single steady-state flux–species coordinate used here.

## Limitations of the study

FSS samples flux–species space uniformly; the implied parameters can land in biophysically implausible regions. The remedy is rejection or importance sampling against a prior on (***k, K***^cat^, ***q***); that choice is situational and outside this paper’s scope. FSS samples fixed points only; oscillatory or transient regimes require sampling along trajectories rather than at steady state, a natural extension via the same machinery. Note that FSS is *orthogonal* to refinements of the sampler itself (quasi-Monte-Carlo schemes, Sobol or lattice sequences, adaptive sampling): all of these can be applied on top of FSS in flux–species space exactly as they are applied in parameter space, and gain the coordinate advantage of FSS. The natural comparisons are therefore not against alternative sampling *distributions* but against alternative sampling *coordinates* (e.g., sampling reduced totals, or sampling in some other algebraic invariant of the network); these we leave to future work.

### Take-home recommendation

Numerical sampling remains essential for contemporary systems biology, but a behavior-discovery workflow should sample the coordinate where behavior is actually read. Flux–species sampling does this directly: draw the fluxes and species first, invert for parameters, and then apply the same stability and phenotype checks already used downstream. The result is not merely a higher hit rate or a better statement of finite-sampling uncertainty; it is a way to recover missed behavior and map it to the dominance conditions and parameter regimes that realise it. We therefore recommend FSS as the default exploratory coordinate for fixed-point biocircuit behaviors, with parameter-space filters used afterward to encode system-specific priors.

## Supplemental information

The Supplemental Information contains the derivations, proofs, controls, and auxiliary figures needed to reproduce or audit the main conclusions. The supplemental items are related to the main figures as follows.

- Table S1, related to Figures 1 and 2: coordinate charts for parameter and flux–species descriptions.
- Figure S1, related to Figure 1: dominance regimes of the sequestration toy.
- Figure S2, related to Figure 2: chain-length interpretation of finite-sample dominance-regime coverage and parameter-side compound inequalities.
- Figure S3, related to Figure 4: two FSS protocols for multistability.
- Figure S4, related to Figures 2 and 3: reading design rules from one sampled state.
- Figure S5, related to Figures 2 and 3: holistic discovery followed by scenario filtering.
- Figure S6, related to Figure 3: post-hoc adaptation filtering under abundance ceilings.
- Figure S7, related to Figure 3: Michaelian complex-estimate mismatch among the recovered adaptation hits.
- Figure S9, related to Figure 3: all-hit comparison of fixed-*σ* and *σK*_*B*1_*/w*_*B*_ setpoint errors among adaptation hits.
- Figure S8, related to Figure 3: a fixed-*σ* JSA-box hit passing our adaptation filter.
- Figure S10, related to Figure 4: dominance-regime analysis of a representative *N* = 10 all-state FSS hit.
- Figure S11, related to Figure 4: asymmetric high-dimensional MultiFate stress test.

