## Supplemental Information for "Flux–species sampling enables holistic exploration of biocircuit behavior"

This supplement is self-contained. The first sections are tutorial: the binding–catalysis formalism and its three running examples (§S1), the dominance-regime picture (§S2), and the coordinate charts (§S3) that explain what FSS computes and why every sampled point is valid. It then proves the completeness guarantee, including the partial-order finite-sample certificate that makes a flux–species null meaningful (§S4, especially §S4.3), derives every local behavior from a sampled state (§S5), records the regime-discovery and scenario-filtering tools (§S6, §S7), and closes with the two case studies (§S8, §S9). Sections, equations, and claims are numbered with an "S" prefix throughout; references to "Theorem 1" and the like without a prefix point to the main text.

### S1 The binding–catalysis framework

This section introduces the objects used by FSS and writes the three running circuits in binding–catalysis form. Following the sequestration toy through this section and the next shows how a flux–species draw is converted into parameters, and why every positive draw is already a valid state of the model. The full theory behind these constructions—existence, uniqueness, well-posedness of the inversion, and the associated geometry—is developed in Liu et al. [3], Xiao [5], Xiao et al. [7]; here we use only the operational consequences. The three circuits recur throughout: their dominance regimes (§S2), charts (§S3), behaviors (§S5), and full case studies (§S8, §S9).

#### S1.1 The layered system

Almost every mechanistic biocircuit model is built from two kinds of reaction, and the framework separates them by timescale. *Binding* reactions—a protein–ligand interaction, monomer dimerisation, or enzyme–substrate association—are fast and reversible, so they sit at equilibrium. *Catalysis* reactions—synthesis, degradation, an enzyme turning substrate into product—are slow and effectively irreversible, and they are what drives the circuit's dynamics. Writing the fast part as algebraic constraints and the slow part as the dynamics gives the differential–algebraic form (1) used throughout [3,5,7]:

$$\tfrac{d}{dt}\boldsymbol{q} = \boldsymbol{S}\boldsymbol{v}, \qquad \boldsymbol{v} = \boldsymbol{K}^{\text{cat}}\boldsymbol{x}, \qquad \boldsymbol{L}\boldsymbol{x} = \boldsymbol{q}, \qquad \boldsymbol{N}\log\boldsymbol{x} = \log\boldsymbol{k}.$$

It is useful to read the system one equation at a time. The state is the vector of $n$ *species* concentrations $\boldsymbol{x} > 0$—every distinct molecular form, free or complexed.

- $\boldsymbol{L}\boldsymbol{x} = \boldsymbol{q}$ (conservation): each *total* $q_i$ is a conserved sum of species—a moiety pool that binding moves among forms but cannot create or destroy. $\boldsymbol{L}$ is a small-integer bookkeeping matrix (which species count toward which pool). There are $d$ totals.
- $\boldsymbol{N}\log\boldsymbol{x} = \log\boldsymbol{k}$ (binding equilibrium): the fast reactions are at equilibrium. Mass action $k_j = \prod_\ell x_\ell^{N_{j\ell}}$ becomes *linear* once you take logarithms—which is why $\log\boldsymbol{x}$ is the natural coordinate. There are $r = n - d$ such relations, one per independent binding reaction, with constants $\boldsymbol{k}$.
- $\boldsymbol{v} = \boldsymbol{K}^{\text{cat}}\boldsymbol{x}$ and $\dot{\boldsymbol{q}} = \boldsymbol{S}\boldsymbol{v}$ (catalysis): the slow *fluxes* $\boldsymbol{v}$ (each a rate constant times its driving species) push the totals around according to the catalysis stoichiometry $\boldsymbol{S}$. A *binding-only* circuit has no catalysis ($m_{\text{cat}} = 0$) and is purely algebraic, fixed by $(\boldsymbol{q}, \boldsymbol{k})$.

For example, in the toy $A + B \rightleftharpoons C$, the two totals are $A_{\text{tot}} = A + C$ and $B_{\text{tot}} = B + C$ (the rows of $\boldsymbol{L}$), and the single binding equilibrium is $K = AB/C$, i.e. $\log A + \log B - \log C = \log K$ (the row of $\boldsymbol{N}$). Given the three species $(A, B, C)$ these are immediate—and that easy direction is precisely what FSS exploits (§S3).

#### S1.2 Two stoichiometric splits and the steady-state manifold

Two simple bookkeeping facts set the dimensions FSS works in. First, *binding* ties the $n$ species down to $d$ independent totals plus $r = n - d$ reaction directions: fixing the totals $\boldsymbol{q}$ and the binding constants $\boldsymbol{k}$ pins the positive species $\boldsymbol{x}$ uniquely for the mass-action binding networks considered here, so $n = r + d$. Second, *catalysis* need not move every total—some combinations stay constant even on the slow timescale. Writing $r_{\text{cat}} = \operatorname{rank} \boldsymbol{S}$ for the number of totals catalysis can actually move and $d_{\text{cat}} = d - r_{\text{cat}}$ for the rest,

$$r_{\text{cat}} := \operatorname{rank} \boldsymbol{S} \ \text{(independent catalytic directions)}, \qquad d_{\text{cat}} := d - r_{\text{cat}} \ \text{(catalysis conservation laws)},$$

a basis $\boldsymbol{W}$ of the left-null-space of $\boldsymbol{S}$ ($\boldsymbol{WS} = 0$) collects the conserved combinations $\boldsymbol{w} = \boldsymbol{Wq}$, the *moieties*: a total enzyme amount, a recycled cofactor pool, a substrate-plus-product sum—quantities fixed for the whole experiment. So the totals split $d = r_{\text{cat}} + d_{\text{cat}}$ into $r_{\text{cat}}$ that *move* (the dynamical directions $\boldsymbol{q}_I$) and $d_{\text{cat}}$ that are *held* (the moieties $\boldsymbol{w}$), exactly mirroring the binding split $n = r + d$[7].

At steady state the nonconserved slow totals stop, $\boldsymbol{Sv} = 0$, so the flux lives in the $(m_{\text{cat}} - r_{\text{cat}})$-dimensional space $\ker \boldsymbol{S}$ and the steady states form a manifold of dimension

$$n + (m_{\text{cat}} - r_{\text{cat}}), \tag{S1}$$

the $n$ species together with the independent fluxes. *This manifold is exactly what FSS samples*—and because $(\boldsymbol{v}, \boldsymbol{x})$ parametrise it directly, every draw is already a steady state, with no equation left to solve. For the toy, $m_{\text{cat}} = 0$ and the manifold is the three species; for the adaptation circuit below ($n = 10$, $m_{\text{cat}} = 4$, $r_{\text{cat}} = 2$) it is $10 + 2 = 12$-dimensional.

#### S1.3 The framework applied to three running examples

We record each circuit's $(\boldsymbol{L}, \boldsymbol{N}, \boldsymbol{S}, \boldsymbol{K}^{\text{cat}})$ once, for reuse.

**(i) Sequestration toy (binding-only).** One reaction $A + B \rightleftharpoons C$, species $\boldsymbol{x} = (A, B, C)$, dissociation constant $K$. With $\boldsymbol{q} = (A_{\text{tot}}, B_{\text{tot}})$ and $\log K = \log \frac{AB}{C}$,

$$\boldsymbol{L} = \begin{array}{c|ccc} & A & B & C \\ \hline A_{\text{tot}} & 1 & & 1 \\ B_{\text{tot}} & & 1 & 1 \end{array}, \qquad \boldsymbol{N} = \begin{array}{c|ccc} & A & B & C \\ \hline K & 1 & 1 & -1 \end{array},$$

and no catalysis ($m_{\text{cat}} = 0$). Here $n = 3$, $d = 2$, $r = 1$; the steady-state manifold (S1) is the three species. The behavior studied is the both-bound ultrasensitive regime, $C/A \geq R$ and $C/B \geq R$.

**(ii) Adaptation circuit (binding–catalysis).** Ten species, ordered

$$\boldsymbol{x} = (I, A, A^*, B, B^*, E, C_1, C_2, C_3, C_4).$$

The four bindings (5) are $I + A \rightleftharpoons C_1$, $A^* + B^* \rightleftharpoons C_2$, $A^* + B \rightleftharpoons C_3$, and $E + B^* \rightleftharpoons C_4$. They give the binding matrix $\boldsymbol{N}$ via $\boldsymbol{N} \log \boldsymbol{x} = \log \boldsymbol{k}$ with $\boldsymbol{k} = (K_{A1}, K_{A2}, K_{B1}, K_{B2})$ (each row one dissociation, as in the toy):

$$\boldsymbol{N} = \begin{array}{c|cccccccccc} & I & A & A^* & B & B^* & E & C_1 & C_2 & C_3 & C_4 \\ \hline K_{A1} & 1 & 1 & & & & & -1 & & & \\ K_{A2} & & & 1 & & 1 & & & -1 & & \\ K_{B1} & & & 1 & 1 & & & & & -1 & \\ K_{B2} & & & & & 1 & 1 & & & & -1 \end{array}$$

The six binding-conserved totals $\boldsymbol{q} = \boldsymbol{L}\boldsymbol{x}$ are the moiety pools $(q_I, q_E, q_A, q_{A^*}, q_B, q_{B^*})$:

$\boldsymbol{L} =$

| | $I$ | $A$ | $A^*$ | $B$ | $B^*$ | $E$ | $C_1$ | $C_2$ | $C_3$ | $C_4$ |
|---|---|---|---|---|---|---|---|---|---|---|
| $q_I$ | 1 | | | | | | 1 | | | |
| $q_E$ | | | | | | 1 | | | | 1 |
| $q_A$ | | 1 | | | | | 1 | | | |
| $q_{A^*}$ | | | 1 | | | | | 1 | 1 | |
| $q_B$ | | | | 1 | | | | | 1 | |
| $q_{B^*}$ | | | | | 1 | | | 1 | | 1 |

so $n = 10$, $r = 4$, $d = 6$. Catalysis carries four fluxes $\boldsymbol{v} = (k_{A1}C_1, k_{A2}C_2, k_{B1}C_3, k_{B2}C_4) = \boldsymbol{K}^{\text{cat}}\boldsymbol{x}$, one nonzero per row of $\boldsymbol{K}^{\text{cat}}$ (column-specific: $\boldsymbol{K}^{\text{cat}}_{1,C_1}{=}k_{A1}$, $\boldsymbol{K}^{\text{cat}}_{2,C_2}{=}k_{A2}$, $\boldsymbol{K}^{\text{cat}}_{3,C_3}{=}k_{B1}$, $\boldsymbol{K}^{\text{cat}}_{4,C_4}{=}k_{B2}$), with catalysis stoichiometry

$\boldsymbol{S} =$

| | $v_1$ | $v_2$ | $v_3$ | $v_4$ |
|---|---|---|---|---|
| $q_A$ | $-1$ | 1 | | |
| $q_{A^*}$ | 1 | $-1$ | | |
| $q_B$ | | | $-1$ | 1 |
| $q_{B^*}$ | | | 1 | $-1$ |

($q_I, q_E$ rows zero).

Hence $\operatorname{rank}\boldsymbol{S} = r_{\text{cat}} = 2$, $\ker\boldsymbol{S} = \{v_1{=}v_2,\ v_3{=}v_4\}$ (dimension 2), and the $d_{\text{cat}} = 4$ conserved moieties are $\boldsymbol{w} = (q_I, q_E, w_A, w_B)$ with $w_A = q_A{+}q_{A^*}$, $w_B = q_B{+}q_{B^*}$. The two nonzero rows of $\boldsymbol{S}\boldsymbol{v}$ are the active-form dynamics $\dot{q_{A^*}} = k_{A1}C_1 - k_{A2}C_2$, $\dot{q_{B^*}} = k_{B1}C_3 - k_{B2}C_4$. A draw of the ten species and the two balanced fluxes ($v_1{=}v_2{=}v_A$, $v_3{=}v_4{=}v_B$) fixes $\boldsymbol{k}$, the totals, and the rates $k_{A1}{=}v_A/C_1$, $k_{A2}{=}v_A/C_2$, $k_{B1}{=}v_B/C_3$, $k_{B2}{=}v_B/C_4$; the studied behavior is free-$A^*$ adaptation (§S8).

**(iii) MultiFate (binding–catalysis), $N = 2$.** The general $N$ repeats this pattern. Five species $\boldsymbol{x} = (A_1, A_2, A_{11}, A_{22}, A_{12})$—two free monomers, two homodimers, one heterodimer. The three dimerisations share $K_d$ and give $\boldsymbol{N}\log\boldsymbol{x} = \log\boldsymbol{k}$ with $\boldsymbol{k} = (K_d, K_d, K_d/2)$ (the factor 2 in $A_{12} = 2A_1A_2/K_d$ absorbed into $\boldsymbol{k}$):

$\boldsymbol{N} =$

| | $A_1$ | $A_2$ | $A_{11}$ | $A_{22}$ | $A_{12}$ |
|---|---|---|---|---|---|
| $A_{11}$ | 2 | | $-1$ | | |
| $A_{22}$ | | 2 | | $-1$ | |
| $A_{12}$ | 1 | 1 | | | $-1$ |

,

$\boldsymbol{L} =$

| | $A_1$ | $A_2$ | $A_{11}$ | $A_{22}$ | $A_{12}$ |
|---|---|---|---|---|---|
| $A_1^{\text{tot}}$ | 1 | | 2 | | 1 |
| $A_2^{\text{tot}}$ | | 1 | | 2 | 1 |

,

the totals being $A_i^{\text{tot}} = A_i + 2A_{ii} + \sum_{j\neq i} A_{ij}$; so $n = 5$, $r = 3$, $d = 2$. Catalysis is the self-activating expression: each total is produced by a saturating (Hill) function of its own homodimer and removed by linear dilution,

$$\dot{A}_i^{\text{tot}} = \underbrace{a + b\,\frac{A_{ii}^n}{1 + A_{ii}^n}}_{\text{production}} - \underbrace{A_i^{\text{tot}}}_{\text{dilution}}, \qquad i = 1, 2,$$

so both totals move ($r_{\text{cat}} = 2$, no conserved moiety, $d_{\text{cat}} = 0$).

The Hill production law can be embedded in the same framework as an effective fast promoter-occupancy readout. For a promoter total $q_D = D + C$, write

$$nA_{ii} + D \rightleftharpoons C, \qquad K_H = \frac{A_{ii}^n D}{C}.$$

With $\boldsymbol{x}_\mathrm{H} = (A_{ii}, D, C)$, this readout has

$$\boldsymbol{N}_\mathrm{H} = \begin{array}{c|ccc} & A_{ii} & D & C \\ \hline K_H & n & 1 & -1 \end{array}, \qquad \boldsymbol{L}_\mathrm{H} = \begin{array}{c|ccc} & A_{ii} & D & C \\ \hline q_{A_{ii}} & 1 & & \\ q_D & & 1 & 1 \end{array}.$$

A physically sequestering promoter would contribute $nC$ to the $A_{ii}$ total, but Zhu et al.'s reduced Hill model treats promoter binding as a non-depleting readout of the homodimer signal, hence $q_{A_{ii}} = A_{ii}$ in this effective module. Solving the readout gives

$$\frac{C}{q_D} = \frac{A_{ii}^n}{K_H + A_{ii}^n}.$$

Thus the model term $bA_{ii}^n/(1 + A_{ii}^n)$ is the catalytic flux $k_\mathrm{cat}C$ after the normalization $K_H = 1$ and $k_\mathrm{cat}q_D = b$. FSS draws the free-monomer levels $A_i$, forms the dimers and totals through $\boldsymbol{N}, \boldsymbol{L}$, and inverts the steady-state balance (production = dilution) for $(a, b)$ (§S9.2).

#### S2 Bioregulatory modes are dominance regimes

This section gives the geometric intuition formalised later in the supplement. A circuit's qualitative behavior is determined by *which species and which fluxes dominate*. Those choices partition coordinate space into dominance regimes, each with a corresponding reduced model. FSS samples the coordinate system in which these regimes are simple chambers. The underlying machinery—dominance profiles, reaction orders, and well-posedness—is developed in Xiao [5], Xiao et al. [6,7]; here we use only the consequences needed for sampling.

**Binding: which species carries each total.** A binding network's behavior at a state is set not by the precise concentrations but by which species carries each conserved total. Writing each total $q_i = \sum_j L_{ij}x_j$ as a competition among its terms, the dominance profile $D_{ij} = L_{ij}x_j / \sum_\ell L_{i\ell}x_\ell$ (2) records the winners; when one term dominates each total ($D_{ij} \approx 1$) the network collapses to a small reduced description—a few monomial relations in $\log \boldsymbol{x}$. This is exactly the move behind the textbook reductions: for an enzyme $E + S \rightleftharpoons C \to E + P$, choosing $S \gg C$ gives Michaelis–Menten, $C \gg S$ (saturation) gives the Hill limit, and $C \gg E$ gives the regime where the enzyme sits mostly bound. Each choice of one dominator per total is a *bioregulatory mode*, or *dominance regime*; the reduced model is not an approximation imposed by hand but the regime's own asymptotic form.

**Toy example: four dominance modes.** The sequestration toy $A + B \rightleftharpoons C$ (§S1.3) makes this concrete (Fig. S1). It has two comparisons—$C$ versus $A$ in the $A$-total, $C$ versus $B$ in the $B$-total—so $2 \times 2 = 4$ modes, one per quadrant of the plane $(\log \frac{C}{A}, \log \frac{C}{B})$: *both free* ($A, B \gg C$, so $A_\mathrm{tot} \approx A$); *$A$-sequestered* ($C \gg A$, so $A_\mathrm{tot} \approx C$ and the free fraction $A/A_\mathrm{tot}$ is small); *$B$-sequestered* ($C \gg B$); and the *both-bound* corner ($A_\mathrm{tot} \approx C \approx B_\mathrm{tot}$). The behavior matched to the ultrasensitivity toy is the both-bound mode with margin, $C/A \geq R$ and $C/B \geq R$: the complex dominates both free pools, so the two totals are both carried by $C$. Its boundaries are the *fixed* lines $A = C$ and $B = C$: in species coordinates they sit at the origin of each comparison and do not move with the binding constant $K$. In parameter coordinates the same regime is seen only after solving the binding equilibrium, where it becomes a diagonal, ill-conditioned condition involving totals and $K$, rather than a direct pair of dominance comparisons.

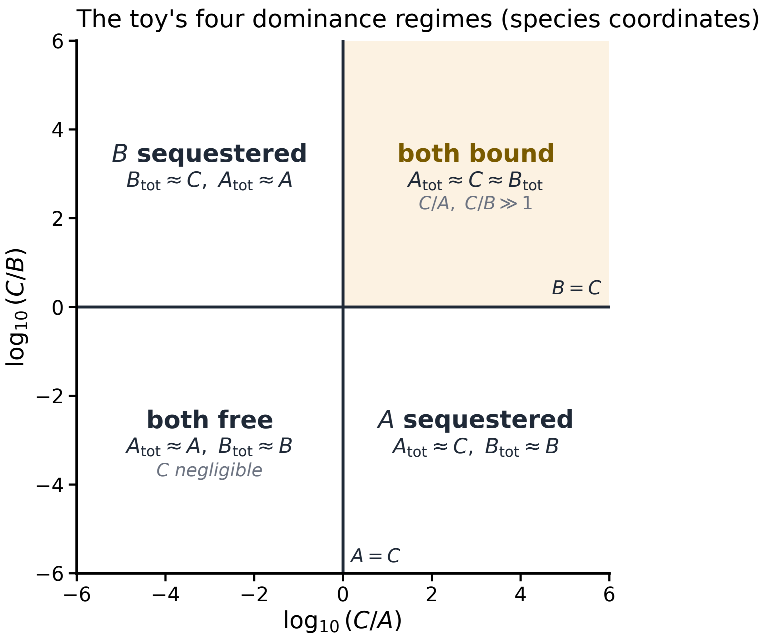

**Figure S1. Dominance regimes of the sequestration toy.** The two comparisons ($C$ vs $A$, $C$ vs $B$) split the species plane into four quadrants—four bioregulatory modes, each with a direct reduced description. The studied behavior (ultrasensitive both-bound dominance, gold) is the mode $C \gg A$ and $C \gg B$. The boundaries $A = C$, $B = C$ are *fixed* in species coordinates, independent of the binding constant $K$; this is why a single box meets every regime (§S4).

**Catalysis: which flux carries each turnover.** With catalysis the same idea applies one layer up. Each independent total's turnover $\dot{q}_{I,i} = \sum_{\text{prod}} v - \sum_{\text{deg}} v$ is a competition among *fluxes*: which flux dominates production and which dominates degradation sets the "birth–death" mode, and balancing the two dominant fluxes gives the steady-state setpoint. Both layers—binding among species, birth–death among fluxes—are first-order comparisons, hence *linear* in $(\log \boldsymbol{x}, \log \boldsymbol{v})$.

Formally, write the slow stoichiometry as

$$\boldsymbol{S} = \boldsymbol{S}^+ - \boldsymbol{S}^-, \qquad S^+_{ij} = \max(S_{ij}, 0), \quad S^-_{ij} = \max(-S_{ij}, 0).$$

Then for each moving total,

$$\dot{q}_{I,i} = \sum_j S^+_{I,ij} v_j - \sum_j S^-_{I,ij} v_j.$$

Flux dominance compares like-unit terms within the production sum $\{S^+_{I,ij} v_j : S^+_{I,ij} > 0\}$ and within the degradation sum $\{S^-_{I,ij} v_j : S^-_{I,ij} > 0\}$, together with the balance between the dominant production and degradation terms at steady state. These are again difference arrangements in $\log \boldsymbol{v}$. The joint binding–catalysis regime is therefore a chamber of a difference arrangement in $\boldsymbol{z} = (\log \boldsymbol{x}, \log \boldsymbol{v}) \in \mathbb{R}^{n+m}$, with the species and flux layers occupying complementary coordinate blocks.

**Dominance regimes as sampling chambers.** Because every comparison is a hyperplane (a log-ratio held above or below a stoichiometric offset), the modes are the *chambers* these hyperplanes cut the flux–species chart into—a tiling of log-space into polyhedra, the four quadrants of Fig. S1 in miniature. Three properties make this the right place to sample: (i) the boundaries are fixed by stoichiometry, never by the rate constants, so the chambers do not move as parameters vary; (ii) each chamber has a simple reduced description—every total a monomial, every turnover a single-term birth–death balance—recovered by reading the dominances off *any* state in it (§S6); and (iii) because the chambers are bounded-offset and scale-free, one finite box meets every one (§S4). A log-uniform draw therefore lands in each mode with room to spare—the whole basis of the completeness guarantee.

### S3 Coordinate charts: parameter space versus flux–species space

The same state is named two ways: the *parameters* a standard pipeline draws, and the *flux–species* coordinates FSS draws. Which variables play which role depends on the kind of problem (Table S1); in all cases the two charts carry the same state and are exchanged by the algebra of (1).

| problem type | flux–species (FSS) | parameters (standard) | determined |
|---|---|---|---|
| binding only | $\boldsymbol{x}$ | $(\boldsymbol{q}, \boldsymbol{k})$ | — |
| binding–catalysis (dynamic) | $(\boldsymbol{v}, \boldsymbol{x})$ | $(\boldsymbol{q}_I, \boldsymbol{w}, \boldsymbol{k}, \boldsymbol{K}^{\text{cat}})$ | — |
| binding–catalysis (steady state) | $(\boldsymbol{v}, \boldsymbol{x}),\ \boldsymbol{v} \in \ker \boldsymbol{S}$ | $(\boldsymbol{w}, \boldsymbol{k}, \boldsymbol{K}^{\text{cat}})$ | nonconserved totals $\boldsymbol{q}_I$ |

**Table S1.** Coordinate charts by problem type. Each row's two charts have the same dimension—$n$ for binding-only, $n + m_{\text{cat}}$ for the dynamic case, $n + (m_{\text{cat}} - r_{\text{cat}})$ at steady state (S1)—and are interchanged by inverting (1): $\boldsymbol{k} = \exp(\boldsymbol{N} \log \boldsymbol{x})$, $\boldsymbol{q} = \boldsymbol{L}\boldsymbol{x}$, and $\boldsymbol{K}^{\text{cat}}$ from $\boldsymbol{v} = \boldsymbol{K}^{\text{cat}}\boldsymbol{x}$. At steady state the balance $\boldsymbol{S}\boldsymbol{K}^{\text{cat}}\boldsymbol{x} = 0$ fixes the nonconserved slow totals $\boldsymbol{q}_I$, so they leave the external parameter list. The three examples of §S1.3 populate the rows: the toy is binding-only, the adaptation and MultiFate circuits are steady-state binding–catalysis.

**Remark S1** (control-knob form)**.** At steady state the parameter side $(\boldsymbol{w}, \boldsymbol{k}, \boldsymbol{K}^{\text{cat}})$ is itself the natural "control-knob" chart: the externally set dials are the conserved amounts $\boldsymbol{w}$ (a total enzyme $q_E$, a regulator pool $w_B$) and the affinities $\boldsymbol{k}$, while the nonconserved totals $\boldsymbol{q}_I$ are whatever the steady-state balance makes them. Setpoints such as the adaptation value $\sigma K_{B1}/w_B$ (§2.4.2) are read in this chart.

#### S3.1 The flux–species coordinate at steady state

With fluxes $\boldsymbol{v} = \boldsymbol{K}^{\text{cat}}\boldsymbol{x}$ ($\boldsymbol{K}^{\text{cat}} \in \mathbb{R}^{m\times n}_{\geq 0}$), the system (1) reads $\dot{\boldsymbol{q}} = \boldsymbol{S}\boldsymbol{v}$, $\boldsymbol{L}\boldsymbol{x} = \boldsymbol{q}$, $\boldsymbol{N} \log \boldsymbol{x} = \log \boldsymbol{k}$. Two facts make $(\boldsymbol{v}, \boldsymbol{x})$ the natural sampling coordinate. First, $\boldsymbol{S}$ may be rank-deficient; a nonnegative $\boldsymbol{W}$ with $\boldsymbol{W}\boldsymbol{S} = 0$ gives conserved pools $\boldsymbol{w} = \boldsymbol{W}\boldsymbol{q}$ (for example a total enzyme or cofactor pool). Splitting the totals into independent and dependent parts $\boldsymbol{q} = (\boldsymbol{q}_I, \boldsymbol{q}_D)$ and writing $\bar{\boldsymbol{L}} = [\boldsymbol{W};\, P_I]\boldsymbol{L}$, a steady state solves

$$\boldsymbol{S}_I\, \boldsymbol{v} = 0, \qquad \boldsymbol{v} = \boldsymbol{K}^{\text{cat}}\boldsymbol{x}, \qquad \bar{\boldsymbol{L}}\, \boldsymbol{x} = (\boldsymbol{w}, \boldsymbol{q}_I), \qquad \boldsymbol{N} \log \boldsymbol{x} = \log \boldsymbol{k}, \tag{S2}$$

and the externally controlled parameters are $(\boldsymbol{w}, \boldsymbol{k}, \boldsymbol{K}^{\text{cat}})$. Second, biological catalytic specificity means each species drives at most one flux column of $\boldsymbol{K}^{\text{cat}}$; an enzyme catalysing several reactions does so through distinct substrate complexes, i.e. distinct species.

Under this specificity the map $(\boldsymbol{v}, \boldsymbol{x}) \mapsto (\boldsymbol{w}, \boldsymbol{k}, \boldsymbol{K}^{\text{cat}})$ is explicit: $\boldsymbol{x}$ gives $\boldsymbol{k} = \exp(\boldsymbol{N} \log \boldsymbol{x})$ and $\boldsymbol{w} = \boldsymbol{W}\boldsymbol{L}\boldsymbol{x}$, and each flux $v_i = \sum_j K^{\text{cat}}_{ij} x_j$ fixes the rate constants of the species driving flux $i$ (uniquely when one species drives the flux, and up to the usual sloppiness among co-drivers otherwise). Binding-only circuits carry no $\boldsymbol{v}$ and FSS reduces to species sampling; direct parameter sampling of $(\boldsymbol{w}, \boldsymbol{k}, \boldsymbol{K}^{\text{cat}})$ provides the comparison.

##### S3.2 Algebraic inversion between charts

The choice of chart matters because the two directions have different computational structure. The map *from* flux–species coordinates *to* parameters is branch-free algebra; the reverse map, required after a parameter draw, is a nonlinear solve and can fail or select a different branch. In the toy, a draw $(A, B, C)$ gives the parameters directly,

$$K = \tfrac{AB}{C}, \qquad A_{\text{tot}} = A + C, \qquad B_{\text{tot}} = B + C,$$

and every positive $(A, B, C)$ yields a valid $(A_{\text{tot}}, B_{\text{tot}}, K)$. The reverse map, drawing $(A_{\text{tot}}, B_{\text{tot}}, K)$ and recovering the species, means solving the quadratic $C^2 - (A_{\text{tot}}+B_{\text{tot}}+K)\,C + A_{\text{tot}}B_{\text{tot}} = 0$ for the complex and then selecting a positive physical root. The two charts represent the same state, but one direction is explicit whereas the other is a potentially branched solve.

With catalysis the asymmetry is even sharper. FSS draws the species *and* the independent fluxes $\boldsymbol{v} \in \ker \boldsymbol{S}$; then $\boldsymbol{k} = \exp(\boldsymbol{N} \log \boldsymbol{x})$, $\boldsymbol{q} = \boldsymbol{L}\boldsymbol{x}$, and the rate constants come from $\boldsymbol{v} = \boldsymbol{K}^{\text{cat}}\boldsymbol{x}$ (one flux per driving species, so each rate is just $v/$[its species]). Because the draw already lies in $\ker \boldsymbol{S}$, the result is a *steady state by construction*: no ODE integration or steady-state solve is required. This is the Fig. 2: where the standard pipeline draws $(\boldsymbol{k}, \boldsymbol{K}^{\text{cat}}, \boldsymbol{q})$ and simulates to steady state, FSS draws $(\boldsymbol{v}, \boldsymbol{x})$ and inverts. For the adaptation circuit, a draw of the ten species and the two balanced fluxes $(v_A, v_B)$ fixes all four dissociation constants, the six totals, and the four catalytic rates $k_{A1} = v_A/C_1,\ k_{A2} = v_A/C_2,\ k_{B1} = v_B/C_3,\ k_{B2} = v_B/C_4$, and the point is a fixed point of the slow dynamics. Thus solver validity is built into the coordinate: FSS evaluates each accepted positive draw directly rather than first searching for a fixed point. The geometric reason to sample this chart is developed in §S2 and §S4; this section only records the algebraic direction that makes each sampled point valid.

#### S4 Completeness of the flux–species box

This section proves the geometric completeness statement behind FSS: a canonical flux–species box intersects every full-dimensional behavior-realizing dominance regime with positive measure, once the box is wide enough for the requested dominance margin. Finite-sample nulls are a consequence of this positive-measure result, not the central object of the section; the sample-budget bound is developed afterwards in §S4.3.

*Step 1: behaviors are read on dominance regimes.* A regime-determined qualitative behavior is true or false on a whole dominance chamber at once (§S2). In the minimal binding toy, for example, the ultrasensitive/sequestered regime is the displayed partial order $C \gg A$ and $C \gg B$. More complex behaviors add flux comparisons and local-stability tests, but the dominance part remains a set of first-order comparisons among species and fluxes.

*Step 2: in flux–species coordinates the chamber arrangement is canonical.* A boundary between two binding dominances is a balance such as $L_{ij}x_j = L_{i\ell}x_\ell$, or $\log x_j - \log x_\ell = \log(L_{i\ell}/L_{ij})$. The offset is set by stoichiometry, not by unknown kinetic constants. With catalysis, the same statement holds for like-unit flux contributions. Thus the dominance chambers form a difference arrangement in log flux–species coordinates. A bounded origin-centred box with an explicit radius meets every full-dimensional chamber; demanding a stronger fold margin simply enlarges the radius by the log of that margin.

*Step 3: positive measure gives a finite-sampling interpretation.* If a realisable behavior occupies a full-dimensional chamber, then its intersection with the declared box has positive volume. An independent log-uniform draw therefore hits it with probability $p > 0$, and $N$ independent draws miss it with probability $(1-p)^N$. Theorem 2 is this positive-measure statement; the chain-length certificate of §S4.3 supplies conservative lower bounds for $p$.

#### S4.1 Formal statements and proofs

The proof proceeds in three steps. Regimes are chambers of a difference arrangement; a box of explicit radius meets every chamber (Lemma S1); that radius sets the quantitative dominance margin (Remark S2); and log-uniform sampling gives a geometrically decaying false-negative probability in the stated scope (Theorem 2). The required polyhedral geometry is from Xiao[5], Xiao et al.[7].

**Modes are chambers of a difference arrangement.** Write $y = \log \boldsymbol{x} \in \mathbb{R}^n$. For a conservation $i$ the competing terms are $\{L_{ij}x_j : L_{ij} > 0\}$, and the dominance boundary between terms $j$ and $\ell$ is $L_{ij}x_j = L_{i\ell}x_\ell$, i.e.

$$y_j - y_\ell \;=\; c_{j\ell}^{(i)}, \qquad c_{j\ell}^{(i)} := \log \tfrac{L_{i\ell}}{L_{ij}}.$$

Each boundary is a hyperplane whose normal is the *difference* $e_j - e_\ell$ and whose offset is bounded by the stoichiometry,

$$\big|c_{j\ell}^{(i)}\big| \;\le\; \beta \;:=\; \max_i \max_{j,\ell:\, L_{ij}, L_{i\ell}>0} \Big|\log \tfrac{L_{i\ell}}{L_{ij}}\Big|.$$

Let $\mathcal{H}$ be the finite arrangement of all such hyperplanes. Its *chambers* (connected components of $\mathbb{R}^n \setminus \bigcup \mathcal{H}$) are exactly the open regions on which every pairwise dominance comparison has a definite sign; on each chamber the dominance profile $D(\boldsymbol{x})$ (2)—and hence the reduced description, the reaction orders, and the sign structure of the Jacobian (3)—is constant. A dominance regime, a choice of one dominating species per conservation, is a union of chambers, so it suffices to reach chambers. Every hyperplane of $\mathcal{H}$ is invariant under the translation $y \mapsto y + t\mathbf{1}$ (a difference $y_j - y_\ell$ ignores $t$), so is every chamber: *which mode a state occupies depends only on ratios of species, never on their overall scale, and never on* $\boldsymbol{k}$ *or* $\boldsymbol{K}^{\mathrm{cat}}$.

**A bounded box meets every mode.**

**Lemma S1** (Completeness radius)**.** *Let $M_\star = \frac{1}{2}(n-1)\beta$. For every $M > M_\star$ the open box $B_M = (-M, M)^n \subset \mathbb{R}^n$ meets the interior of every chamber of $\mathcal{H}$—hence every dominance regime of the network.*

*Proof.* Fix a chamber $P$, cut out by one strict inequality per hyperplane. Rewriting each as a lower bound on a difference, $P = \{y : y_{u_k} - y_{v_k} > w_k,\ k = 1, \dots, m\}$ with $|w_k| \le \beta$. Form the weighted digraph $G$ on $\{1, \dots, n\}$ with an edge $v_k \to u_k$ of weight $w_k$. Since $P \neq \varnothing$ the system is feasible, so $G$ carries no cycle of positive weight (the feasibility criterion for difference constraints). Set

$$y_j^\star \;=\; \max\{\mathrm{wt}(\pi) : \pi \text{ a simple path of } G \text{ ending at } j\},$$

including the empty path (weight 0); then $0 \le y_j^\star \le (n-1)\beta$, since a simple path uses at most $n-1$ edges of weight $\le \beta$. For any edge $v \to u$ of weight $w$, extending an optimal path to $v$ gives $y_u^\star \ge y_v^\star + w$ (if appending closes a cycle, that cycle has weight $\le 0$, so the bound still holds), whence $y^\star \in \overline{P}$ with spread $\max_j y_j^\star - \min_j y_j^\star \le (n-1)\beta$; translating along $\mathbf{1}$ centres it to $\|y^\star\|_\infty \le M_\star$. Finally $P$ is open, convex, and nonempty, so every neighbourhood of $y^\star \in \overline{P}$ contains a point of $P$; for any $M > M_\star$ such an interior point lies in $B_M$. □

For *unit stoichiometry* ($L_{ij} \in \{0,1\}$) one has $\beta = 0$ and $M_\star = 0$: every mode's interior reaches arbitrarily close to the origin, and *any* box captures all of them. Non-unit coefficients (e.g. dimerisation, $L = 2$) only enlarge $M_\star$ to $\frac{1}{2}(n-1)\log(\text{max coefficient ratio})$, a small number of log-units even for large networks. The box size is therefore set not by *which* modes exist—all are intersected—but by how much quantitative *strength* the behavior demands.

**Quantitative strength sets the radius.**

**Remark S2.** A behavior demanding a tunable dominance strength enlarges $M_\star$ additively by the log of that strength, and no more. In the sequestration toy, the both-bound margin $C/A \geq R$ and $C/B \geq R$ is $y_A - y_C \leq -\log R$ and $y_B - y_C \leq -\log R$. Thus a box with $M > M_\star + \log R$ contains the strengthened dominance chamber. The box grows only like $\log R$: a stronger threshold means a more complete dominance relation, not a new coordinate system.

Lemma S1 together with this strength margin is exactly Theorem 1.

**Finite samples from positive-measure regimes.** Call a behavior $X$ *regime-determined* if its truth value is constant on each dominance regime; ultrasensitivity beyond a fixed fold, a prescribed adaptation regime, a co-stable state pattern, and the Jacobian sign conditions of §2.2 are all of this form. Say $X$ is *realisable* if some full-dimensional regime satisfies it.

*Proof of Theorem 2.* Fix $M > M_\star$ and draw log-coordinate points i.i.d. uniformly from $B_M$. By Lemma S1 a realising regime has interior meeting $B_M$; that intersection is open and nonempty, hence of positive volume, so the per-draw hit probability $p = \text{vol}\{y \in B_M : X\}/\text{vol}(B_M)$ is strictly positive. The draws are independent, so all $N$ miss with probability $(1-p)^N \to 0$, and the complementary event—a realisable $X$ yielding an unbounded run of empty samples—has probability zero. Thus, within the declared full-dimensional regime class and box, repeated FSS sampling recovers any realisable regime-determined behavior with probability tending to one. □

This is the finite-sample consequence of positive measure: if the behavior is full-dimensional and regime-determined inside the declared box, the false-negative rate decays geometrically in the sample size. The practical role of this statement is constructive. A behavior that parameter sampling misses because the corresponding regime is thin, shifted, or ill-conditioned in parameter coordinates can still be recovered by FSS in the flux–species box where that regime has ordinary volume. It is not a proof that every possible transient behavior is absent, nor does it cover exact-equality slices of zero measure. Liu et al.[3] establish the companion fact that every dominance chamber has positive measure; Lemma S1 adds that a single bounded box of explicit radius already intersects every chamber, so the guarantee survives the move from an idealised unbounded sample to a practical finite one.

**Scope.** Three assumptions delimit the result. (i) *Regime-determined behaviors.* The statement covers properties fixed by the dominance regime and the Jacobian sign-structure—the qualitative phenotypes this paper samples for. A property that *also* requires the slow catalysis rates to lie in a range is handled by adjoining $\log \boldsymbol{K}^{\text{cat}}$ to the coordinates: rate comparisons are again differences, so $\mathcal{H}$ and Lemma S1 extend verbatim with $n$ replaced by $n$ plus the number of rate coordinates. (ii) *Full-dimensional regimes.* A behavior confined to a measure-zero boundary (an exact-equality setpoint) has $p = 0$ in *both* coordinate systems; Section S8.5 is such a case, and FSS correctly returns nothing. (iii) *Totality of the inversion.* Every $y \in B_M$ yields a genuine state with $\log \boldsymbol{k} = \boldsymbol{N}y$ and $\boldsymbol{q} = \boldsymbol{L}e^y$ all positive, so no draw is lost to infeasibility; the sample size $N$ in Theorem 2 is the *effective* one, unlike parameter sampling, where solving for $\boldsymbol{x}$ may return no positive branch.

#### S4.2 Formal chain-length geometry

This subsection gives the formal object behind the certificate below. Definition S1 defines chain length as the smallest box a regime forces; Proposition S1 connects it to the completeness radius; Lemma S2 and Theorem S1 show why the species and flux layers remain separated in $(\boldsymbol{v}, \boldsymbol{x})$. The parameter-coordinate failure mode is separated into §S4.4. The polyhedral geometry is Xiao[5], Xiao et al.[7].

**Chain length, canonically.** How wide a box a regime needs is set not by the *number* of its conditions but by how *nested* they are, and the right measure of nesting is forced on us by *units*. Every validity condition compares two quantities of the *same physical dimension* (a concentration with a concentration, a flux with a flux), so written as

$$\langle c, \boldsymbol{z} \rangle \geq \delta \qquad (\delta = \log \text{fold}) \tag{S3}$$

the exponent vector $c$ *balances within every fundamental dimension*: the ratio is genuinely dimensionless. Hence the regime cone $\{\boldsymbol{z} : \langle c_i, \boldsymbol{z} \rangle \geq \delta \ \forall i\}$ is invariant under the *dilation group*—one independent log-shift of the coordinates carrying each dimension (here, concentration and time)—because changing a unit only rescales within its dimension. (This also forbids a spurious chain such as $M \gg M^2 \gg M^3$, which would need a dimension-violating, degree-changing comparison.)

**Definition S1** (Chain length)**.** Group the coordinates by physical dimension. The *chain length* of a regime $R$ in coordinate system $Z$ is the minimal coordinate spread its $\delta$-margin conditions (S3) force, measured *modulo the dilation group*—the smallest, over the cone, of the largest per-dimension span:

$$\ell_Z(R) \;=\; \frac{1}{\delta} \min_{\langle c_i, \boldsymbol{z} \rangle \geq \delta \ \forall i} \ \max_{\text{dimensions } g} \Big( \max_{j \in g} z_j - \min_{j \in g} z_j \Big), \tag{S4}$$

a linear program. Within each dimension the optimiser sorts coordinates into scale bands—a graded poset; $\ell_Z(R)$ is the height of the tallest. Measuring modulo the dilation group is what makes $\ell_Z$ a property of the regime and coordinate system, not of the arbitrary choice of concentration and time units.

Because (S4) is an LP optimum, $\ell_Z(R)$ is *exact*: no longer chain can be hiding, since a longer forced ordering would enlarge the spread. Shift-centring the optimiser gives $\ell_Z(R)\,\delta = 2M_\star(R)$—**chain length is exactly twice the per-regime completeness radius** of Lemma S1, the rigorous form of the heuristic $M_\star = \frac{1}{2}\beta\ell$.

**Proposition S1** (Chain length sets the box)**.** *A box of half-width $M > \frac{1}{2}\delta\,\ell_Z(R)$ meets $R$, and none smaller does; hence $M_\star(Z) = \frac{1}{2}\delta\ \max_R \ell_Z(R)$ and the probability a uniform $N$-draw misses $R$ grows with $\ell_Z(R)$. Chain length is therefore the sampling-difficulty currency for a declared dominance regime: longer nested orders require a wider box and give smaller conservative hit probabilities.*

**Examples of chain length.** Definition S1 is terse, so we unpack it on small regimes (read "$\gg$" as "exceeds by a factor $\geq 10^\delta$"). In each, $\ell$ is the minimal coordinate spread (S4) in units of $\delta$, after grouping coordinates by dimension; we state the minimising assignment and read off the spread.

*(i) One dominance.* $R = \{x_1 \gg x_2\}$ has the single constraint $z_1 - z_2 \geq \delta$, so the spread $z_1 - z_2$ is minimised at $\delta$: $\ell = 1$, bands $\{x_2\} \prec \{x_1\}$.

*(ii) A dominator over a crowd.* $R = \{x_1 \gg x_2,\ x_1 \gg x_3\}$: take $z_2{=}z_3{=}0,\ z_1{=}\delta$; spread $\delta$, $\ell = 1$, bands $\{x_2, x_3\} \prec \{x_1\}$. *Many conditions need not make a long chain*—a star is one step.

*(iii) A transitive chain.* $R = \{x_1 \gg x_2,\ x_2 \gg x_3\}$ forces $z_1 - z_3 \geq 2\delta$; spread $2\delta$, $\ell = 2$, bands $\{x_3\} \prec \{x_2\} \prec \{x_1\}$. *Nesting is what costs*: a link forced to sit *between* two others adds a band.

*(iv) A product comparison.* $R = \{x_1x_2 \gg x_3x_4\}$, one balanced condition $z_1 + z_2 - z_3 - z_4 \geq \delta$: split the burden, $z_1{=}z_2{=}\frac{\delta}{4}$, $z_3{=}z_4{=}-\frac{\delta}{4}$; spread $\frac{\delta}{2}$, $\ell = \frac{1}{2}$. *A product dominance is cheaper than a pairwise one*—no single coordinate carries the whole gap, so $\ell$ can fall below 1. This is why the multi-term conditions typical of parameter coordinates do not, on their own, make chains long.

The integer $l$ used in the operational certificate of §S4.3 is the displayed-partial-order special case of $\ell_Z$: it is what (S4) returns when all regime constraints are simple pairwise comparisons in one dimension group. Product inequalities can have fractional $\ell_Z$, so they are handled by this formal definition rather than by the displayed-order probability bound.

**Decoupling and completeness in $(\boldsymbol{v}, \boldsymbol{x})$.** The reason $(\boldsymbol{v}, \boldsymbol{x})$ keeps chains short is structural: the two dominance layers never share a coordinate.

**Lemma S2** (Layer decoupling)**.** *In $\boldsymbol{z} = (\log \boldsymbol{x}, \log \boldsymbol{v})$ every binding hyperplane involves only $\log \boldsymbol{x}$ and every birth–death hyperplane only $\log \boldsymbol{v}$. Hence each joint regime is a product chamber $P_{\boldsymbol{x}} \times P_{\boldsymbol{v}}$ with*

$$\ell_{(\boldsymbol{v},\boldsymbol{x})}(R) = \max\{\ell_{\boldsymbol{x}}(P_{\boldsymbol{x}}),\, \ell_{\boldsymbol{v}}(P_{\boldsymbol{v}})\},$$

*and, the offsets being log-ratios of entries of $\boldsymbol{L}$ and $\boldsymbol{S}$ alone, $\ell_{(\boldsymbol{v},\boldsymbol{x})}(R)$ depends only on the topology $(\boldsymbol{L}, \boldsymbol{S})$ and the regime, never on the constants $(\boldsymbol{w}, \boldsymbol{k}, \boldsymbol{K}^{\mathrm{cat}})$.*

*Proof.* A binding condition compares $L_{ij}x_j$ with $L_{i\ell}x_\ell$, i.e. $\log x_j - \log x_\ell = \log(L_{i\ell}/L_{ij})$; a birth–death condition compares $|S_{ij}|v_j$ with $|S_{i\ell}|v_\ell$. The two families occupy complementary coordinate blocks of $\boldsymbol{z}$ and share no variable, so no transitive chain passes from an $\boldsymbol{x}$-monomial to a $\boldsymbol{v}$-monomial: the longest chain is the longer of the two layers'. The offsets are fixed by $\boldsymbol{L}, \boldsymbol{S}$; the constants enter only the *values* a draw takes, not which chamber contains it. □

**Theorem S1** (Completeness and finite-sample consequence in $(\boldsymbol{v}, \boldsymbol{x})$)**.** *Let $\beta$ bound the binding offsets $|\log(L_{i\ell}/L_{ij})|$ and birth–death offsets $|\log(|S_{i\ell}|/|S_{ij}|)|$. For every $M > \frac{1}{2}\beta \max_R \ell_{(\boldsymbol{v},\boldsymbol{x})}(R)$ the box $(-M, M)^{n+m}$ in $\boldsymbol{z}$ meets the interior of every regime, so flux–species sampling realises each regime-determined behavior with positive probability and its $N$-draw false-negative rate decays geometrically.*

*Proof.* The binding and birth–death conditions are difference-type in $\boldsymbol{z} \in \mathbb{R}^{n+m}$, so the joint regimes are chambers of a difference arrangement and Lemma S1 applies: each chamber's interior meets a box of half-width $\frac{1}{2}\beta$ times the longest constraint path, which by Lemma S2 is $\ell_{(\boldsymbol{v},\boldsymbol{x})}(R)$. The geometric decay is Theorem 2 applied to $\boldsymbol{z}$, valid because regime-determined behaviors are constant on these chambers. □

#### S4.3 Operational finite-sample certificate for dominance regimes

This subsection turns the positive-measure guarantee and the chain-length definition above into a finite-sample certificate. Dominance requirements are displayed partial orders among log flux/species coordinates. For this operational certificate, $l$ is the integer height of the displayed partial order: a star with one dominator and many dominated terms still has $l = 1$, whereas a nested ladder $X \gg Y \gg Z$ has $l = 2$. If $X_1, \ldots, X_n$ are independent log-coordinates sampled uniformly from an interval of width $m$, and each displayed edge $u \succ v$ means

$$X_u \geq X_v + \delta,$$

then the worst-case single-draw probability among all displayed partial orders on $n$ variables with longest chain length $l$ is

$$p_l^\star(n,\rho) = \frac{(q_l!)^{\,l-r_l}\,((q_l+1)!)^{r_l}}{n!}\,[1-(l-1)\rho]_+^{\,n}, \qquad n = q_l l + r_l, \quad 0 \le r_l < l, \quad \rho = \frac{\delta}{m}.$$

For $d$ independent samples, a fixed target with single-draw probability $p$ is missed with probability $(1-p)^d$. Thus the 99% required per-draw probability is

$$p_{\mathrm{req}}(d) = 1 - 0.01^{1/d},$$

and the maximum covered chain length is

$$l_{\max}(d) = \max\{l : p_l^\star(n,\rho) \ge p_{\mathrm{req}}(d)\}.$$

If a behavior contains an independent species partial order and an independent flux partial order, the conservative lower bound multiplies:

$$p_{\mathrm{hit}}^\star = p_{l_x}^\star(n_x,\rho_x)\, p_{l_v}^\star(n_v,\rho_v).$$

This product is the explicit lower bound for the $p$ used in Theorem 2. For adaptation-like steady-state phenotypes, the procedure is: read the dominance conditions and local-response sign/precision conditions at the sampled fixed point, reduce the dominance part to species and flux partial orders, and report which chain lengths the sample budget covers. This is a conservative certificate for the pairwise displayed-order case; product inequalities and parameter-coordinate comparisons are interpreted by the formal $\ell_Z$ definition in §S4.2.

#### S4.4 Coordinate inflation after eliminating fluxes

The point of the flux–species chart is not that parameter coordinates are always numerically longer. It is that $(\boldsymbol{v},\boldsymbol{x})$ keeps the displayed dominance order literal: species dominances are species comparisons, and birth–death dominances are flux comparisons. After eliminating the fluxes through $\boldsymbol{v} = \boldsymbol{K}^{\mathrm{cat}}\boldsymbol{x}$, an atomic flux comparison becomes a product inequality among rates and species. If that flux dominance has to overcome an opposite species dominance on the same complexes, the parameter-side regime contains a longer nested chain.

The minimal pattern is a dilute-but-fast route: $x_A \ll x_B$ in concentration but $v_A \gg v_B$ in flux. In $(\boldsymbol{v},\boldsymbol{x})$ these are two independent single-step comparisons, one in the concentration block and one in the flux block. Eliminating the flux ($v = kx$) gives $k_A x_A \gg k_B x_B$; together with $x_B \gg x_A$, this forces

$$\frac{k_A}{k_B} \gg \frac{x_B}{x_A} \gg 1, \tag{S5}$$

a two-step ladder in the parameter chart.

**Proposition S2** (Chain inflation in conflicting flux regimes)**.** *Eliminating the fluxes via $\boldsymbol{v} = \boldsymbol{K}^{\mathrm{cat}}\boldsymbol{x}$ turns an atomic flux dominance into a product inequality involving rates and species. When that flux dominance must overcome an opposite binding dominance on shared species, the parameter-side displayed order contains a strictly longer nested chain than the flux–species order.*

*Sketch.* A birth–death condition $v_A \gg v_B$, atomic in $\boldsymbol{v}$, becomes $\sum_\ell K_{A\ell} x_\ell \gg \sum_\ell K_{B\ell} x_\ell$ after flux elimination. On a dominance regime its leading terms reduce this to a product comparison

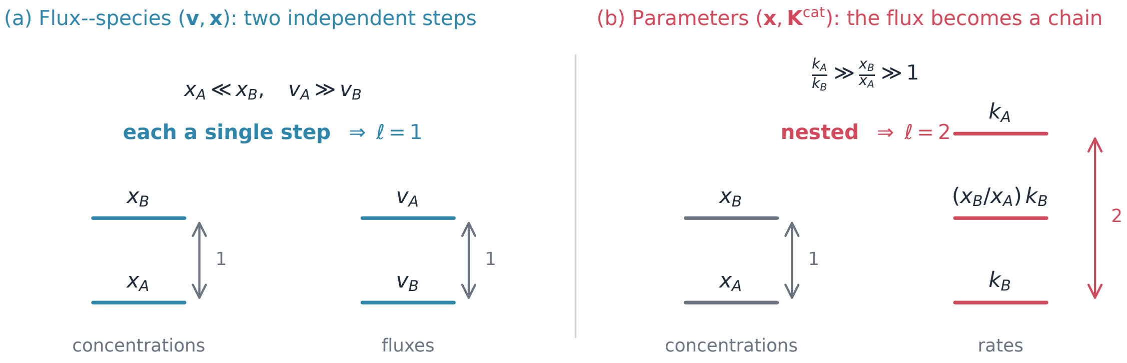

**Figure S2. Chain length, and how parameter coordinates can inflate it.** A regime needs $x_A \ll x_B$ (concentrations) and $v_A \gg v_B$ (fluxes). (a) In flux–species coordinates the two conditions are independent single steps, one per dimension group: chain length $\ell = 1$, so a one-decade box reaches the regime. (b) Eliminating the flux ($v = kx$) makes $v_A \gg v_B$ a product that must overcome the concentration gap, nesting the rate group into a two-step ladder $k_A/k_B \gg x_B/x_A \gg 1$: chain length $\ell = 2$, a wider box and $\sim\ell$-fold rarer hits.

$K_{A,a}x_a \gg K_{B,b}x_b$. If the same regime also fixes $x_b \gg x_a$, the rate ratio must compensate for the species gap:

$$K_{A,a}/K_{B,b} \gg x_b/x_a \gg 1.$$

This is the length-2 chain of (S5), whereas $(\boldsymbol{v}, \boldsymbol{x})$ had two independent single links in separate coordinate blocks. □

**Remark S3** (a structural bound, and what is open)**.** By Lemma S2, $\ell_{(\boldsymbol{v},\boldsymbol{x})}(R)$ is a finite invariant of the topology and regime, independent of rate constants. Parameter coordinates need not always have a longer numerical chain length, because product inequalities can distribute a gap across several parameters. Their weakness is different: the regime is no longer a literal partial order among the variables being sampled. The adaptation regime has short chains on both sides but compound parameter inequalities and a stricter setpoint criterion; the competing-flux toy below has a true chain-length inflation.

The dilute-but-fast pattern above is sufficient for the chain-length point: eliminating flux variables can turn separate dominance links in $(\boldsymbol{v}, \boldsymbol{x})$ into nested parameter inequalities.

### S5 Behaviors from flux–species

A sampled flux–species state $(\boldsymbol{v}, \boldsymbol{x})$ is a steady state of the closed dynamics by construction (§S1), so every *local* behavior of the circuit there—stability, steady-state sensitivity, ultrasensitivity, adaptation precision, input-responsiveness, and (across states) multistability—is fixed by the Jacobian of the slow dynamics. This section is the self-contained derivation: from a draw we get the controller's response to perturbations by one linear solve (§S5.2), compose it into the slow Jacobian in closed form (§S5.3), and then read each phenotype off that Jacobian (§S5.4–§S5.7). No ODE is integrated and no parameter is perturbed; every formula is evaluated once per sampled point. The same machinery serves both case-study samplers, so their comparison is exact.

##### S5.1 A sampled state is a fixed point: what is computable

A draw fixes the species $\boldsymbol{x}$ and the independent fluxes $\boldsymbol{v} \in \ker \boldsymbol{S}$, and hence (§S1) the binding constants $\boldsymbol{k} = \exp(\boldsymbol{N} \log \boldsymbol{x})$, the totals $\boldsymbol{q} = \boldsymbol{L}\boldsymbol{x}$, and a catalysis matrix with $\boldsymbol{v} = \boldsymbol{K}^{\text{cat}}\boldsymbol{x}$. Because $\boldsymbol{S}\boldsymbol{v} = 0$, the nonconserved slow totals are balanced and $\boldsymbol{x}$ is a steady state of $\dot{\boldsymbol{q}} = \boldsymbol{S}\boldsymbol{v}$ on the binding manifold $\{\boldsymbol{L}\boldsymbol{x} = \boldsymbol{q},\ \boldsymbol{N} \log \boldsymbol{x} = \log \boldsymbol{k}\}$. Two derivatives govern all local behavior: how the equilibrium species move when a total is perturbed—the fast binding layer (§S5.2)—and how the totals themselves evolve—the slow catalysis layer (§S5.3).

##### S5.2 Species response to total perturbations

On the binding manifold $\boldsymbol{L}\boldsymbol{x} = \boldsymbol{q}$ and $\boldsymbol{N} \log \boldsymbol{x} = \log \boldsymbol{k}$. Differentiating both, with the binding constants $\boldsymbol{k}$ held fixed,

$$\boldsymbol{L}\, d\boldsymbol{x} = d\boldsymbol{q}, \qquad \boldsymbol{N}\, \text{diag}(1/\boldsymbol{x})\, d\boldsymbol{x} = 0,$$

and stacking the blocks gives the $n \times n$ matrix and the controller response

$$\boldsymbol{A}_{\boldsymbol{x}} = \begin{bmatrix} \boldsymbol{L} \\ \boldsymbol{N}\, \text{diag}(1/\boldsymbol{x}) \end{bmatrix}, \qquad \frac{\partial \boldsymbol{x}}{\partial q_j}\Big|_{\boldsymbol{k}} = \boldsymbol{A}_{\boldsymbol{x}}^{-1} e_j.$$

$\boldsymbol{A}_{\boldsymbol{x}}$ is invertible wherever the binding network is nondegenerate (the rows of $\boldsymbol{L}$ and of $\boldsymbol{N}\,\text{diag}(1/\boldsymbol{x})$ together span $\mathbb{R}^n$), which holds in the interior of every dominance regime. This one solve returns the full $n \times d$ response $\partial \boldsymbol{x}/\partial \boldsymbol{q}|_{\boldsymbol{k}}$; it is exact, integration-free, and the only linear algebra the readouts below require.

##### S5.3 The slow Jacobian

The slow dynamics move the independent totals, $\dot{\boldsymbol{q}}^{\text{cat}} = \boldsymbol{S}\boldsymbol{v} = \boldsymbol{S}\boldsymbol{K}^{\text{cat}}\boldsymbol{x}$. Composing the catalytic dependence with the controller response of §S5.2 gives the slow Jacobian in closed form,

$$\boldsymbol{J} = \boldsymbol{S}\, \boldsymbol{K}^{\text{cat}}\, \frac{\partial \boldsymbol{x}}{\partial \boldsymbol{q}^{\text{cat}}}\Big|_{\boldsymbol{k}},$$

which is (3) of the main text. Every entry comes from $\boldsymbol{x}$ and the fixed topology $(\boldsymbol{L}, \boldsymbol{N}, \boldsymbol{S}, \boldsymbol{K}^{\text{cat}})$—no rate is perturbed, no trajectory simulated.

##### S5.4 Stability

The sampled state is linearly stable iff every eigenvalue of $\boldsymbol{J}$ has negative real part. This single eigenvalue test is the only stability filter used—identically for both case studies and for both sampling methods (flux–species and parameter)—so the methods are compared under the same stability criterion. All phenotype scores below are evaluated only after this filter.

##### S5.5 Steady-state sensitivity and adaptation precision

Treat an input as one of the totals, $q_{\text{in}} \in \boldsymbol{q}$ (for the adaptation circuit $q_{\text{in}} = q_I$). By the implicit function theorem on the balanced slow system $\boldsymbol{F}(\boldsymbol{q}^{\text{cat}}; q_{\text{in}}) = \boldsymbol{S}\boldsymbol{K}^{\text{cat}}\boldsymbol{x} = 0$, the steady total responds as

$$\frac{d(\boldsymbol{q}^{\text{cat}})^{\text{ss}}}{dq_{\text{in}}} = -\boldsymbol{J}^{-1} \frac{\partial \boldsymbol{F}}{\partial q_{\text{in}}},$$

and the chain rule through $\partial \boldsymbol{x}/\partial \boldsymbol{q}|_{\boldsymbol{k}}$ (§S5.2) propagates it to any free species. For an output $y$ (e.g. free $A^*$) the single number that grades the phenotype is the *logarithmic gain*

$$g_y = \frac{d \log y^{\mathrm{ss}}}{d \log q_{\mathrm{in}}}. \tag{S6}$$

*Adaptation precision* is its reciprocal, $\mathcal{P}_y = |g_y|^{-1}$, large when the output is invariant to the input; the main numerical comparison calls a state adapting (in the Ma et al.[4] sense) when $\mathcal{P}_{A^*} \geq 100$.

##### S5.6 Ultrasensitivity

The same gain (S6), read on a titration instead of held invariant, measures ultrasensitivity: $|g_y| > 1$ sustained over a fold of input is a switch-like, super-linear response (a local Hill coefficient above one), and $|g_y| \leq 1$ is graded. Precision and ultrasensitivity are thus two readings of one quantity—a large $|g_y|$ on the actuator is the ultrasensitive switch of Fig. 1d, a small $|g_y|$ on the regulated output is the adaptation of §S5.5. For a binding-only observable the gain needs no Jacobian at all: it is read directly from the controller response $\partial \boldsymbol{x}/\partial \boldsymbol{q}|_{\boldsymbol{k}}$, which is why the switch-like toy of Fig. 1d is exact algebra. The sign of $|g_y| - 1$ is constant on each dominance chamber, so ultrasensitivity is regime-determined (§S4) and flux–species sampling maps the ultrasensitive modes completely.

##### S5.7 Input-responsiveness

High precision alone is not adaptation: an output that never moves is invariant but functionally unresponsive, so we also require a transient response to the input. Linearise the slow system around the fixed point, with the input a scalar deviation $u$ from baseline,

$$\frac{d\boldsymbol{q}^{\mathrm{cat}}}{dt} = \boldsymbol{J}\, \boldsymbol{q}^{\mathrm{cat}} + \boldsymbol{b}\, u, \qquad \boldsymbol{b} = \frac{\partial \boldsymbol{F}}{\partial q_{\mathrm{in}}},$$

and write the output deviation (free $A^*$) as the linear functional $y(t) = \boldsymbol{c}^\top \boldsymbol{q}^{\mathrm{cat}}(t)$ with $\boldsymbol{c}^\top = (\partial A^*/\partial \boldsymbol{q}^{\mathrm{cat}})|_{\boldsymbol{k}}$ from §S5.2. With zero initial deviation,

$$y(t) = \int_0^t \boldsymbol{c}^\top e^{\boldsymbol{J}(t-s)} \boldsymbol{b}\, u(s)\, ds,$$

so the response is governed by the scalar impulse response $h(t) = \boldsymbol{c}^\top e^{\boldsymbol{J}t} \boldsymbol{b}$. The output is structurally responsive iff some Markov coefficient $\boldsymbol{c}^\top \boldsymbol{J}^k \boldsymbol{b}$ is nonzero; by Cayley–Hamilton it suffices to check $k < d_{\mathrm{cat}}$. For the example adaptation network $d_{\mathrm{cat}} = 2$, so the structural test is $\max(|\boldsymbol{c}^\top \boldsymbol{b}|, |\boldsymbol{c}^\top \boldsymbol{J} \boldsymbol{b}|) > 0$ above numerical noise. The numerical criterion adds an amplitude floor: for a unit log-input step ($\delta \log q_{\mathrm{in}} = 1$) the stable linear step response of $\delta \log A^*(t)$ is evaluated and

$$G_A = \max_{t \geq 0} |\delta \log A^*(t)|,$$

with the sample called input-responsive only when $G_A \geq 0.3$—a tenfold input producing at least a $10^{0.3}$-fold output excursion. $G_A$ uses the matrix exponential of the $2 \times 2$ Jacobian, not a nonlinear simulation.

##### S5.8 Multistability

Multistability asks a different question of a circuit than adaptation: not how one fixed point responds, but how *many* target states are simultaneously stable fixed points of a *single* parameter set. With up to $2^N$ candidate patterns (MultiFate-$N$, §2.5), the design target is a parameter vector at which a prescribed family of states co-exists. Flux–species sampling supplies the states directly, in the chart where they are well-conditioned; two protocols turn that into a multistability search, and we use the second.

**Method A: per-state inversion and parameter closeness.** For each target state $s$ in the desired family, sample a flux–species point $(\boldsymbol{v}, \boldsymbol{x})_s$ consistent with that state's dominance pattern and invert it (§S1) to a parameter vector $\theta_s = (\boldsymbol{w}, \boldsymbol{k}, \boldsymbol{K}^{\text{cat}})_s$. A single circuit realises the whole family only if the $\theta_s$ *coincide*. Since the states are drawn independently, their inverted parameters generally differ, and one reads multistability from the *closeness* of the cloud $\{\theta_s\}$: a region where all targeted states map to nearly the same $\theta$ (within a tolerance) is a candidate multistable circuit. This is fast but heuristic—the verdict depends on the tolerance, and exact coincidence is measure-zero.

**Method B: one consistent parameter set, recompute the rest.** To answer the direct multistability question, the parameter set must be fixed first. We sample one reference state's $(\boldsymbol{v}, \boldsymbol{x})$ and invert it to a *single* $\theta$. Holding $\theta$ fixed, for every other target pattern $s$, solve the circuit's fixed-point equations at $\theta$ for the state carrying that pattern—a Newton solve on the algebraic system (1)—obtaining $\boldsymbol{x}_s(\theta)$ or finding none. Each recovered fixed point is then tested for linear stability by the Jacobian of §S5.4. The number of target patterns realised as stable fixed points *at the same* $\theta$ is the multistability of that circuit. Because $\theta$ is single and consistent by construction, this is exactly the quantity meant by "multistable"—no closeness tolerance enters—and it reuses the same Newton-and-Jacobian verifier as parameter-space sampling, so the two methods are compared with the same verifier. We use Method B throughout.

**The MultiFate instances.** Both case-study analyses are Method B. *Symmetric* MultiFate (§S9.2) exploits permutation symmetry to collapse "recompute every pattern" to one representative per ON-count $M$, with the single inversion the $2 \times 2$ solve for $(a, b)$ from an ON and an OFF free-monomer level; accepted representatives are multiplied by $\binom{N}{M}$. *Asymmetric* MultiFate (§S9.5) is Method B in full $2N$ dimensions: draw an ON and an OFF monomer level per factor, invert *per factor* to $(a_i, b_i)$ through $N$ independent $2 \times 2$ solves—one consistent $\theta$—then count co-stable patterns with the shared $N$-dimensional verifier. In both, the per-state monomer levels are sampled in the complete species chart (§S4). The symmetric case shows the clean all-state recovery result used in Fig. 4; the asymmetric full-$2N$-dimensional stress test in §S9.5 records how that comparison changes when the species window and the per-factor Zhu box are both broadened.

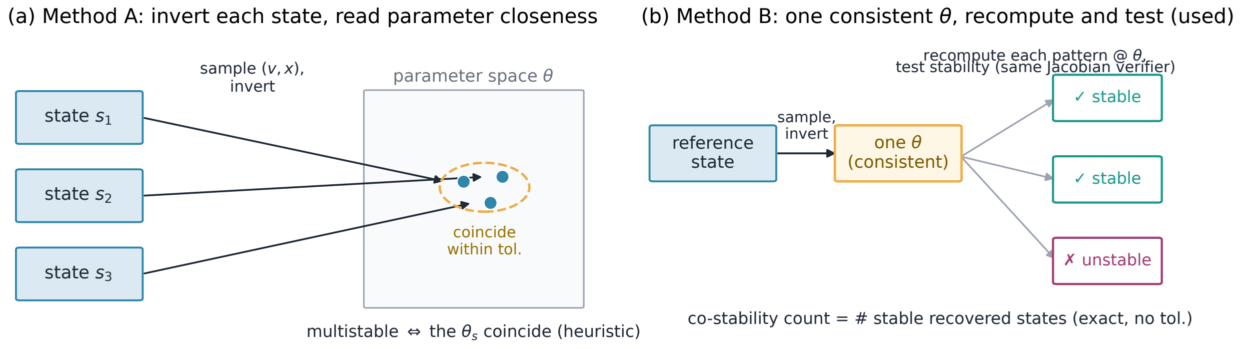

**Figure S3. Two flux–species protocols for multistability.** (a) *Method A*: each target state is sampled in its own flux–species coordinates and inverted to parameters $\theta_s$; multistability is read from the *closeness* of the inverted cloud, and needs a tolerance. (b) *Method B* (used here): one state is sampled and inverted to a single consistent $\theta$; every other target pattern is then *recomputed* as a fixed point at that $\theta$ and tested for stability, giving an exact co-stability count with the same Newton-and-Jacobian verifier as parameter sampling.

### S6 Regime discovery from a sampled point

Sampling returns a point; regime discovery converts that point into the *regime* that supports the behavior: the dominance pattern, the reduced birth–death system, and the symbolic inequalities that define the design rule. This step turns flux–species sampling from point finding into mechanism discovery. From one hit we recover systematic conditions, because a regime-determined behavior is constant on the whole dominance chamber (§S4).

#### S6.1 Reading the dominances off the point

Take a sampled $(\boldsymbol{v}, \boldsymbol{x})$ that realises the behavior. Two sets of quantities are read from the point (Fig. S4):

1. *Binding (static).* For each conserved total $q_i = \sum_j L_{ij} x_j$, find the dominant term $L_{i\,j^*(i)} x_{j^*(i)}$ at the point. The total is then carried by one species, a monomial relation $q_i \approx L_{i\,j^*} x_{j^*}$, and the strict inequalities $\{L_{i\,j^*} x_{j^*} \gg L_{i\ell} x_\ell\}_{\ell \neq j^*}$ cut out the binding chamber (§S4.1).

2. *Birth–death (dynamic).* For each independent total's balance $\dot{q}_{I,i} = \sum_{\text{prod}} S\, v - \sum_{\text{deg}} |S|\, v$, find the dominant production flux and dominant degradation flux. The balance reduces to a single-term-each ODE $\dot{q}_{I,i} \approx v_{\text{prod}^*} - v_{\text{deg}^*}$, and the inequalities selecting them cut out the birth–death chamber (§S4.2).

#### S6.2 The reduced system and the design rule

Substituting the binding monomials into the fluxes $\boldsymbol{v} = \boldsymbol{K}^{\text{cat}} \boldsymbol{x}$ closes the reduction. Read *statically*, setting $v_{\text{prod}^*} = v_{\text{deg}^*}$ gives a monomial fixed-point equation whose solution is the behavior's closed-form setpoint. Read *dynamically*, the one-term-each ODE is the simple birth–death system whose linearisation is the dominant part of the Jacobian (3). Either way, the inequalities collected above are the *conditions* under which the behavior provably holds: one sampled hit yields not a point but a full-dimensional region and its defining rule. That is regime discovery—the systematic complement to sampling.

##### S6.3 Toy instance

*Toy (sequestration).* A hit in the chamber $C \gg A$ and $C \gg B$ gives $A_{\mathrm{tot}} \approx C \approx B_{\mathrm{tot}}$; the rule for strength is $C/A \geq R$ and $C/B \geq R$ (§S4.1)—two like-unit dominances, discovered from one point. The same discovered chamber also proves the local ultrasensitivity of the toy. Fix $A_{\mathrm{tot}}$ and $K$, titrate $B_{\mathrm{tot}}$, and use

$$A_{\mathrm{tot}} = A + C, \qquad B_{\mathrm{tot}} = B + C, \qquad K = \frac{AB}{C}.$$

Implicit differentiation gives

$$g_A = \frac{d \log A}{d \log B_{\mathrm{tot}}}\Big|_{A_{\mathrm{tot}},K} = -\frac{B_{\mathrm{tot}} C}{AB + AC + BC}. \tag{S7}$$

Now write the two free fractions as

$$\epsilon_A = \frac{A}{A_{\mathrm{tot}}}, \qquad \epsilon_B = \frac{B}{B_{\mathrm{tot}}}.$$

Because $A_{\mathrm{tot}} = A + C$ and $B_{\mathrm{tot}} = B + C$, the exact gain becomes

$$|g_A| = \frac{1 - \epsilon_A}{\epsilon_A + \epsilon_B - \epsilon_A \epsilon_B}. \tag{S8}$$

In the $C$-dominated chamber sampled by FSS, $\epsilon_A \ll 1$ and $\epsilon_B \ll 1$, so

$$|g_A| \simeq \frac{1}{\epsilon_A + \epsilon_B} \gg 1.$$

Thus the ultrasensitive response is not an additional hidden condition: it is the local derivative consequence of the same $C$-dominant regime. Quantitatively, if $C/A \geq R$ and $C/B \geq R$, then $\epsilon_A, \epsilon_B \leq 1/(R+1)$ and hence $|g_A| \geq R/2$.

The adaptation and MultiFate instances use the same readout, but their mechanism-specific reductions are kept with their case-study sections: adaptation in §S8.4, and MultiFate in §S9.2.

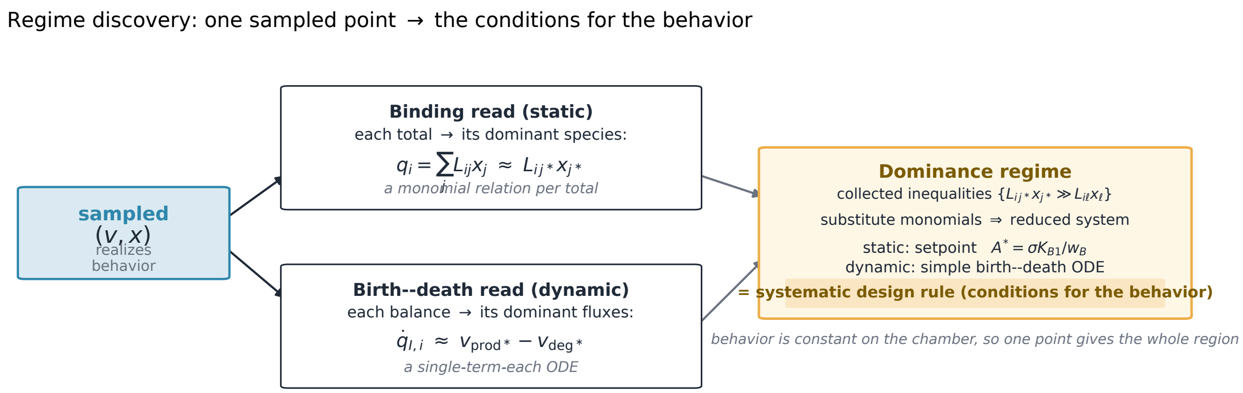

**Figure S4. Regime discovery: one sampled point to a design rule.** A sampled flux–species state is read twice—each conserved total to its dominant species (binding, a monomial relation) and each independent total's turnover to its dominant production and degradation fluxes (birth–death, a single-term-each ODE). The collected strict inequalities are the dominance regime; substituting the monomials closes the reduced system to the behavior's setpoint (static) or simple birth–death ODE (dynamic). Because the behavior is constant on the chamber (§S4), the inequalities are the systematic condition under which it holds, recovered from the single point.

### S7 Two-stage FSS: discovery followed by scenario filtering

The experiments in this section make precise what FSS does and does not claim. The key point is that FSS is a *discovery* tool: from one sampling run in a canonical species range it establishes that a behavior exists and maps the parameter regime that realises it. Whether a discovered regime is "biophysical" is then a separate, scenario-dependent question answered by ordinary post-processing of the same samples. The setpoint-specific adaptation controls are folded into the adaptation case study (§S8); the high-dimensional scaling test into the MultiFate case study (§S9).

FSS deliberately splits exploration into two stages (Fig. S5). *Stage 1—holistic discovery*: one log-uniform sampling run of the canonical box establishes whether a behavior exists and maps the regime that realises it. The completeness guarantee of §S4 is what makes this discovery run broad; finite-sample nulls then inherit a scoped interpretation as a consequence. This stage is a property of the circuit *topology*—it depends on no scenario, and is computed once. *Stage 2—data processing*: any scenario constraint (an abundance ceiling, an affinity window, a specific setpoint) is a direct filter on the *same* sample. Scenarios change; the Stage-1 map does not, so a single holistic exploration is reused for every downstream question. Both case studies are worked instances of exactly this split.

A recurring objection to FSS is that the canonical range $\boldsymbol{x} \in [10^{-3}, 10^{6}]$ implies parameters that may be implausible for a given system. This conflates two steps that FSS deliberately separates. "Biophysical" is not a universal constraint: the plausible range for a dissociation constant or a total abundance shifts by orders of magnitude between a small molecule and a macromolecule, between bacteria and mammalian cells, and between starvation and nutrient-rich growth. There is therefore no single prior to impose *a priori*. FSS instead does discovery first—mapping the regime wherever it lives—after which any scenario constraint is applied as a direct filter to the sampled points. Figure S6 shows this for the adaptation case study. FSS discovers the free-$A^*$ adaptation regime across many decades of $(\log_{10} K_{B1}, \log_{10} w_B)$, obeying the structural requirement $w_B \gg K_{B1}$ of regime (S9). Applying an abundance ceiling $W_{\max}$ to the *same* samples then reads off how many

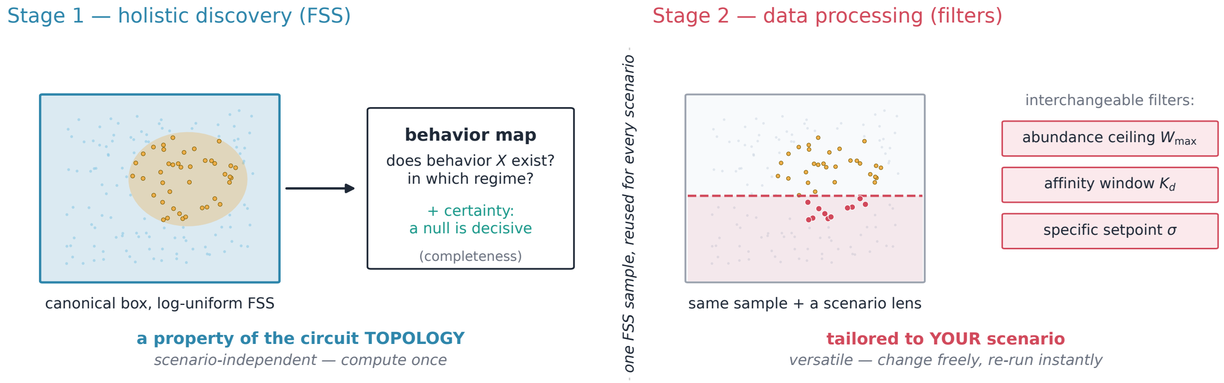

**Figure S5. The two stages of FSS.** *Stage 1 (holistic discovery):* a single log-uniform sampling run of the canonical box maps where a behavior lives (§S4)—a property of the circuit topology, computed once. *Stage 2 (data processing):* scenario filters (an abundance ceiling $W_{\max}$, an affinity window, a specific setpoint) are interchangeable lenses on the same sample. The Stage-1 map is invariant to the scenario; only the filters change.

adaptation designs survive for a given scenario: 0.026%, 0.114%, and 0.319% of raw draws at $W_{\max} = 10^4, 10^5, 10^6$ respectively (generous affinity window $K_d \in [10^{-6}, 10^6]$ applied throughout). One FSS run yields the whole filtering curve; a parameter-box study yields a single point—the one box it chose, here the JSA box at 0.113%—and must re-pick and re-run a box for every new scenario. The pipeline "sample once in the canonical range, then data-process" is the practical point: no up-front prior-guessing, and the same samples serve every downstream plausibility question.

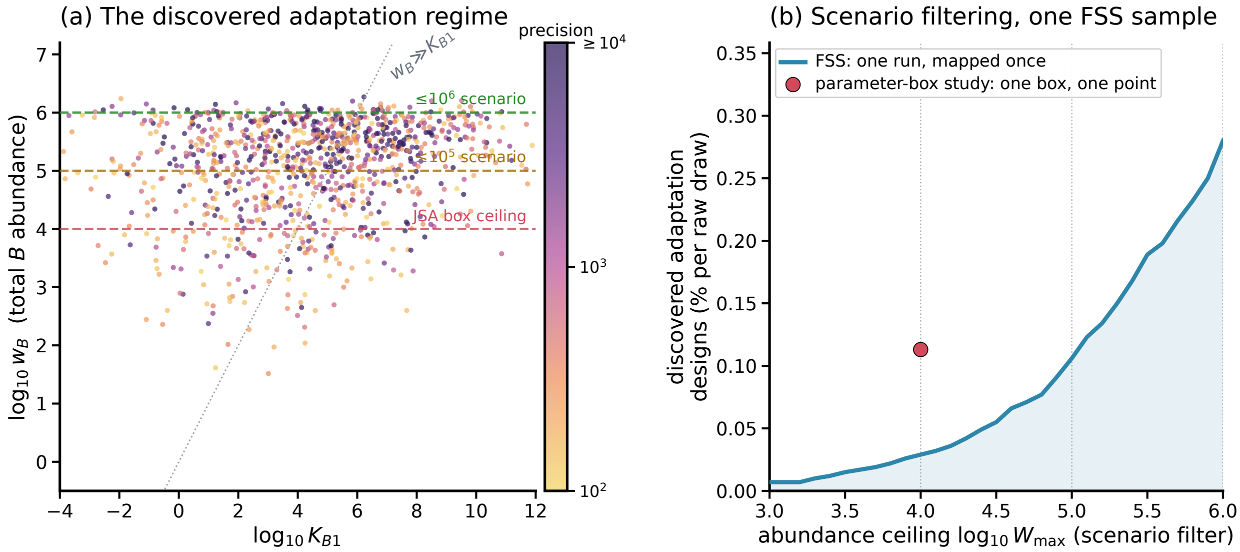

**Figure S6. Discovery, then post-hoc scenario filtering (adaptation).** (a) FSS discovers the free-$A^*$ adaptation regime across ~14 decades of $K_{B1}$ and obeys $w_B \gg K_{B1}$; example scenario abundance ceilings are horizontal lenses on the *same* discovered cloud. (b) Number of discovered adaptation designs surviving an abundance ceiling $W_{\max}$: one FSS run yields the entire curve, whereas a parameter-box study yields one point (its chosen box). Colour in (a) is clipped $\log_{10} P_{A^*}$.

### S8 The adaptation case study

This section is organized as a case study rather than as a collection of diagnostics: first the explicit-complex model, then the Michaelian reduction used by Ma et al., then the sampling and filtering protocol, then the mechanism that parameter sampling missed, and finally why the Jeynes-Smith–Araujo screen returned a negative result.

#### S8.1 Model setup

The adaptation example uses the competitive-binding enzymatic feedback motif summarised in §S1.3. The fixed species vector is

$$\boldsymbol{x} = (I, A, A^*, B, B^*, E, C_1, C_2, C_3, C_4),$$

with binding reactions

$$I + A \rightleftharpoons C_1, \qquad A^* + B^* \rightleftharpoons C_2, \qquad A^* + B \rightleftharpoons C_3, \qquad E + B^* \rightleftharpoons C_4.$$

The active-form totals are

$$q_{A^*} = A^* + C_2 + C_3, \qquad q_{B^*} = B^* + C_2 + C_4,$$

and the two dynamic balances are

$$\dot{q_{A^*}} = k_{A1}C_1 - k_{A2}C_2, \qquad \dot{q_{B^*}} = k_{B1}C_3 - k_{B2}C_4.$$

The input is the total $q_I = I + C_1$, and the scored output is free $A^*$. Thus the case study asks whether free $A^*$ can remain input-invariant after a perturbation of $q_I$, while retaining a non-trivial response in the actuator branch.

#### S8.2 Ma et al.'s Michaelian reduction and adaptation setpoint

Ma et al.'s analysis treats the same feedback logic through a Michaelian reduction: enzyme–substrate complexes are assumed negligible compared with the free substrate pools. In that reduction the $B$-arm balance loses the explicit complex carriers, so the steady-state $q_{B^*}$ equation becomes

$$0 = k_{B1}A^* - k_{B2}q_E.$$

Consequently the reduced model predicts the familiar input-invariant setpoint

$$A^* = \frac{k_{B2}q_E}{k_{B1}} \equiv \sigma.$$

This is how the Michaelian description finds adaptation in this motif: the output level is determined by $B$-arm catalytic constants and the enzyme total, not by the input total $q_I$.

The rest of the case study keeps this Michaelian target as a reference point, but it does not assume that explicit complexes must adapt at this same value. The explicit-complex routes are derived after the sampling protocol, where they can be compared with the recovered hits.

##### S8.3 Sampling method and filtering standard

The adaptation comparison is numerical: neither the sampling code nor the plotted hit mask uses dominance classifications, BNC labels, or regime-interior points. For *parameter-space sampling* we replicated Jeynes-Smith & Araujo's protocol: $10^5$ log-uniform draws from the JSA sampling range (7); for each, Newton on the algebraic flux-balance equations $k_{A1}C_1 = k_{A2}C_2$ and $k_{B1}C_3 = k_{B2}C_4$ (under binding quasi-equilibrium) located the fixed point. For *FSS sampling* we drew all ten species concentrations $(I, A, A^*, B, B^*, E, C_1, C_2, C_3, C_4)$ log-uniformly from the physical concentration range $[10^{-3}, 10^6]$ (sub-nanomolar to high-millimolar; a broad canonical range), computed $\boldsymbol{k}$ from $\boldsymbol{N}\log\boldsymbol{x} = \log\boldsymbol{k}$ and the six totals from $\boldsymbol{L}\boldsymbol{x}$, drew the two balanced catalytic fluxes $(v_A, v_B)$ log-uniformly from $[10^{-3}, 10^6]$, and *derived* all four catalysis rates $k_{A1} = v_A/C_1$, $k_{A2} = v_A/C_2$, $k_{B1} = v_B/C_3$, $k_{B2} = v_B/C_4$ so that $\boldsymbol{x}$ is a fixed point by construction. The analytic Jacobian (3) filtered unstable samples; the implicit function theorem applied to the same Jacobian gave $d\log A^*/d\log q_I$ analytically; the impulse-response criterion §S5.7 gave responsiveness. Adaptation precision is $|d\log A^*/d\log q_I|^{-1}$, called adaptation in the Ma et al. [4] sense when $\geq 100$ for the main finite numerical comparison.

**Scoring.** Each stable sample is graded by the two closed-form Jacobian readouts of §S5: the steady-state log-gain $g_{A^*}$, giving adaptation precision $\mathcal{P}_{A^*} = |g_{A^*}|^{-1}$ (§S5.5), and the step-response amplitude $G_A$, giving input responsiveness (§S5.7). A sample counts as an adaptation hit when $\mathcal{P}_{A^*} \geq 100$ and $G_A \geq 0.3$. Both are evaluated from the sampled state through the analytic Jacobian (3), with no nonlinear simulation.

**Setpoint-specific filters.** Setpoint-specific FSS uses the same cache with an additional filter, not a second sampler. After keeping valid, stable, responsive adaptation hits, we compute $\sigma = k_{B2}q_E/k_{B1}$ for each draw and filter by the log relative error $|\log_{10}(A^*/\sigma)|$. Candidates that are meant to reproduce the literal Jeynes-Smith–Araujo test are then rerun through the ten-input `araujo_rpa` sampling run. This is why FSS can ask either Ma's any-setpoint question or the stricter fixed-$\sigma$ question on the same flux–species sample, while keeping clear which question is full-dimensional and which imposes an extra setpoint constraint. The JSA-box analogue of this stricter question is dissected explicitly in §S8.5, where fixed-$\sigma$ behavior is shown to be an additional equality coincidence rather than the broad adaptation mechanism.

##### S8.4 Adaptation mechanism missed by parameter sampling

**The recovered adaptation is non-Michaelian.** The main text uses the mechanistic negligible-complex test: every explicit complex should be small compared with the free substrate species retained by the Michaelian reduction. For the adaptation network, this means checking the four complex/free ratios

$$\frac{C_1}{A}, \qquad \frac{C_2 + C_3}{A^*}, \qquad \frac{C_3}{B}, \qquad \frac{C_2 + C_4}{B^*}.$$

A conservative 10% negligible-complex bound would require all four ratios to be below 0.1. Across the 1027 FSS adaptation hits, only 2 satisfy this bound, and the median of the largest ratio across the four comparisons is $10^{4.87}$ (Fig. 3e). Thus the adapting states do not merely sit outside a convenient parameter box; they sit outside the assumption that makes the Michaelian reduction valid.

As a companion diagnostic, compare the complexes implied by the Michaelian closure with the real complexes in the explicit competitive-binding state. Using

$$C_1^{\mathrm{MM}} = \frac{q_I q_A}{K_{A1} + q_A}, \quad C_2^{\mathrm{MM}} = \frac{q_{B^*} q_{A^*}}{K_{A2} + q_{A^*}},$$
$$C_3^{\mathrm{MM}} = \frac{q_{A^*} q_B}{K_{B1} + q_B}, \quad C_4^{\mathrm{MM}} = \frac{q_E q_{B^*}}{K_{B2} + q_{B^*}},$$

where $q_A = A + C_1$, $q_{A^*} = A^* + C_2 + C_3$, $q_B = B + C_3$, and $q_{B^*} = B^* + C_2 + C_4$, define

$$R_{\mathrm{MM}} = \max_{i=1,\ldots,4} \frac{|C_i^{\mathrm{MM}} - C_i|}{C_i}.$$

Across the 1027 FSS adaptation hits, $\log_{10} R_{\mathrm{MM}}$ has median 3.43; only 11 hits have worst-complex error below 10%. Thus the Michaelian formulas estimate the complexes themselves incorrectly. Figure S7 separates this complex-estimate diagnostic from the sampling-geometry controls: it asks only whether the explicit-complex fixed points are actually close to their Michaelian closure.

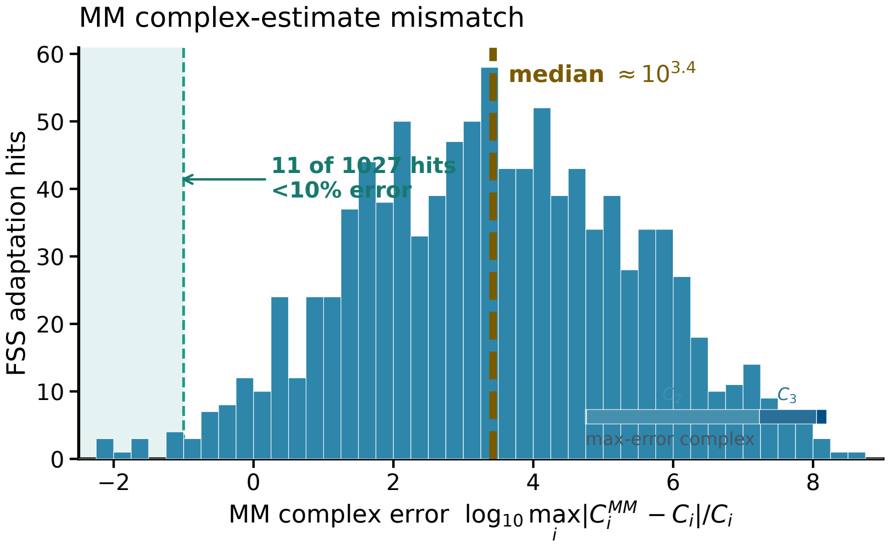

**Figure S7. Michaelian complex estimates fail on the recovered adaptation hits.** For each FSS adaptation hit, the plot shows $R_{\mathrm{MM}} = \max_i |C_i^{\mathrm{MM}} - C_i|/C_i$, the largest relative error between the Michaelian complex estimate and the explicit-complex steady state. Only 11/1027 hits have worst-complex error below 10%, so the recovered adaptation is not captured by the Michaelian complex closure.

**Dominance routes.** The unconstrained FSS hits expose two dominant routes for free-$A^*$ adaptation, and the missed route appears after splitting the free-$E$ branch by the dominant carrier of $w_A$. At a winner-take-all dominance readout, 933/1027 FSS hits are $C_4$-dominated in $q_E = E + C_4$. The remaining 94/1027 are free-$E$-dominated, of which 59 also have $w_A \approx C_2$. In the reconstructed JSA-box hits from the matched $10^5$-draw parameter sample, the corresponding counts are 109 for $q_E \approx C_4$, 0 for $q_E \approx E; w_A \approx C_2$, and 3 for $q_E \approx E$ with other $w_A$ carriers. Thus the parameter box does not merely under-sample the free-$E$ branch; it misses the $C_2$-buffered free-$E$ branch shown in Fig. 3f.

The $C_4$-dominated route is the direct complex-complete counterpart of Ma's intuition. A simple reference chamber is

$$q_{A^*} \approx C_3, \quad q_{B^*} \approx B^*, \quad w_A \approx A, \quad w_B \approx B, \quad q_I \approx C_1, \quad q_E \approx C_4. \tag{S9}$$

The exact $B$-arm balance then gives

$$k_{B1}C_3 = k_{B2}C_4, \qquad C_3 = \frac{A^*B}{K_{B1}}, \qquad A^* = \frac{K_{B1}k_{B2}C_4}{k_{B1}B}.$$

If $C_4 \approx q_E$, then

$$A^* \approx \frac{K_{B1}k_{B2}q_E}{k_{B1}B} = \sigma\,\frac{K_{B1}}{B}.$$

Adaptation in this route therefore requires the free-$B$ denominator to be insensitive to input. The transparent subcase $B \approx w_B$ gives

$$A^* \approx \sigma\,\frac{K_{B1}}{w_B}. \tag{S10}$$

More generally, the denominator can be whichever residual of the $w_B$-total is buffered in the particular dominance chamber.

The free-$E$-dominated route is a distinct mechanism. Here $C_4 = EB^*/K_{B2} \approx q_E B^*/K_{B2}$, so

$$A^* \approx \frac{K_{B1}k_{B2}q_E}{k_{B1}K_{B2}}\frac{B^*}{B}.$$

When $C_2$ dominates the active pools, the equilibrium $C_2 = A^*B^*/K_{A2}$ gives

$$(A^*)^2 \approx \frac{K_{A2}K_{B1}k_{B2}q_E C_2}{K_{B2}k_{B1}B}.$$

Thus the route becomes input-invariant when the ratio $C_2/B$ is buffered. The clean subcase $C_2 \approx w_A$ and $B \approx w_B$ yields

$$A^* \approx \left(\frac{K_{A2}K_{B1}k_{B2}q_E w_A}{K_{B2}k_{B1}w_B}\right)^{1/2}.$$

In the FSS free-$E$ hits, the median $C_2/w_B$ is 0.356 and the median $B/w_B$ is 0.377, showing that this route genuinely uses a $C_2$-rich active pool. In the matched parameter-box hits, the free-$E$ route has median $C_2/w_B = 4.9 \times 10^{-4}$ and $B/w_B = 0.998$; it therefore samples a thin, mostly free-$B$ corner rather than the $C_2$-buffered mechanism. This is the mechanism-level reason the JSA box can contain some adaptation hits while still missing the dominant species-space route.

The parameter-space thinness follows from the same dominance readout. The $A$-arm flux balance gives

$$C_1 = \rho_A C_2, \qquad \rho_A = \frac{k_{A2}}{k_{A1}}.$$

The free-$E$, $C_2$-buffered route has $C_2 \approx w_A$. A visible input response additionally requires that the input not be almost entirely free, so a minimal order-one condition is $I \lesssim C_1$. Since $q_I = I + C_1$, this gives

$$C_1 \lesssim q_I \lesssim 2C_1.$$

Substituting $C_1 = \rho_A C_2$ and $C_2 \approx w_A$ yields the strip

$$1 \lesssim \frac{q_I}{\rho_A w_A} \lesssim 2, \qquad \rho_A = \frac{k_{A2}}{k_{A1}},$$

which is Equation (9) in the main text. In log space this strip has width $\log_{10} 2 \simeq 0.301$ along the coordinated variable $\log_{10} q_I - \log_{10} w_A - \log_{10} \rho_A$. In flux–species coordinates, by contrast, the same condition is an open dominance chamber: $q_E \approx E$, $w_A \approx C_2$, and the input arm remains responsive.

##### S8.5 Setpoint-specific filtering in the Jeynes-Smith–Araujo screen

The released Jeynes-Smith–Araujo code tests a setpoint-specific statement: whether free $A^*$ remains near the Michaelian value $\sigma = k_{B2}q_E/k_{B1}$ while the input is varied. The conclusion stated from that screen is stronger, namely that free $A^*$ does not adapt in the explicit-complex circuit. The FSS hits above separate these two statements: free $A^*$ adapts, but most hits are not fixed-$\sigma$ hits.

**JSA-box search with the same filter used in this work.** To confirm that the published negative is a consequence of a narrow setpoint-specific screen, we ported Jeynes-Smith and Araujo's sampling box and fixed-$\sigma$ criterion from their released code (`RandSamplingCC.m`) into our pipeline. We then imposed the same adaptation filter used throughout this work: stability, nontrivial free $A^*$, precision $\mathcal{P}_{A^*} \geq 100$, and input responsiveness $G_A \geq 0.3$. Finally, for the diagnostic fixed-$\sigma$ subset, we required free $A^*$ to remain within 2% of $\sigma = k_{B2}q_E/k_{B1}$ over a tenfold down–up input range. In a deterministic JSA-box search capped at $5 \times 10^6$ draws, the first point satisfying both our filter and this fixed-$\sigma$ trace appeared only after approximately $2 \times 10^6$ draws. Thus a fixed-$\sigma$ hit that is also responsive is present in the box, but it is a rare target rather than the broad adaptation mechanism.

**A fixed-$\sigma$ hit that passed the filter.** The first such point is informative because a genuine fixed-$\sigma$ hit might seem to revive Ma's Michaelian setpoint as the explanatory mechanism. It does not. The parameter set is

$$\begin{aligned}(K_{A1}, K_{A2}, K_{B1}, K_{B2}) &= (1.535,\ 2.997\times10^2,\ 5.978\times10^3,\ 2.599\times10^{-3}),\\ (k_{A1}, k_{A2}, k_{B1}, k_{B2}) &= (1.057,\ 3.941\times10^2,\ 1.043\times10^{-2},\ 6.452\times10^{-2}),\\ (w_A, w_B, q_E, q_I) &= (4.062\times10^3,\ 6.006\times10^3,\ 15.245,\ 189.178).\end{aligned}$$

At its baseline steady state, $(q_{A^*}, q_{B^*}) = (190.223, 17.319)$, and the explicit species are

$$\begin{aligned}(I, A, A^*, B, B^*, E) &= (7.885\times10^{-2},\ 3.683\times10^3,\ 95.520,\ 5.895\times10^3,\ 1.591,\ 2.486\times10^{-2}),\\ (C_1, C_2, C_3, C_4) &= (189.099,\ 0.507,\ 94.196,\ 15.220).\end{aligned}$$

For this point $\sigma = 94.350$, $A^*/\sigma = 1.012$, $\mathcal{P}_{A^*} = 496.7$, and $G_A = 0.774$. Across $q_I/q_I^0 \in [0.1, 10]$, the largest fixed-$\sigma$ relative error is 1.72%, while the actuator $B^*$ changes by a factor of about 98 (Fig. S8). The point is therefore a genuine adapting state by our filter, but it is still non-Michaelian by the same MM-complex diagnostic: the largest relative error between the Michaelian complex estimate and the real complex is $R_{\mathrm{MM}} = 12.26$ ($\log_{10} R_{\mathrm{MM}} = 1.09$). Its dominance regime,

$$q_I \approx C_1, \quad q_E \approx C_4, \quad w_B \approx B, \quad q_{A^*} \approx A^* + C_3, \quad q_{B^*} \approx C_4,$$

is not the broad regime (S9) with only the input arm changed: $q_{A^*}$ is shared by two comparable terms, $A^* = 95.520$ and $C_3 = 94.196$, rather than being dominated by $C_3$ alone. Nevertheless, the algebra still shows why the hit is close to $\sigma$. The $q_E$ dominance gives $C_4 \approx q_E$; substituting this into the $B$-arm flux balance gives

$$k_{B1}C_3 = k_{B2}C_4 \quad \Rightarrow \quad C_3 \approx \frac{k_{B2}q_E}{k_{B1}} = \sigma.$$

Binding equilibrium then gives the explicit free-output expression

$$C_3 = \frac{A^*B}{K_{B1}} \quad \Rightarrow \quad A^* \approx \sigma\,\frac{K_{B1}}{B}.$$

Thus this point lands on the literal Michaelian value only after an additional parameter/species coincidence: $K_{B1}/B \approx 1$. Numerically, $K_{B1} = 5978 \approx B = 5895$, so $A^*/C_3 = K_{B1}/B = 1.014$ and $A^*/\sigma = 1.012$. Equivalently, once $C_4 \approx q_E$ makes $C_3 \approx \sigma$, the hit passes because $A^*$ happens also to be nearly equal to $C_3$. The total form $\sigma K_{B1}/w_B$ of (S10) is only one approximation to one dominance route, valid when $w_B$ is dominated by free $B$; here it holds strongly ($B/w_B = 0.981$). The strict-$\sigma$ property is therefore an extra equality slice inside an adapting neighborhood, not a full-dimensional dominance regime.

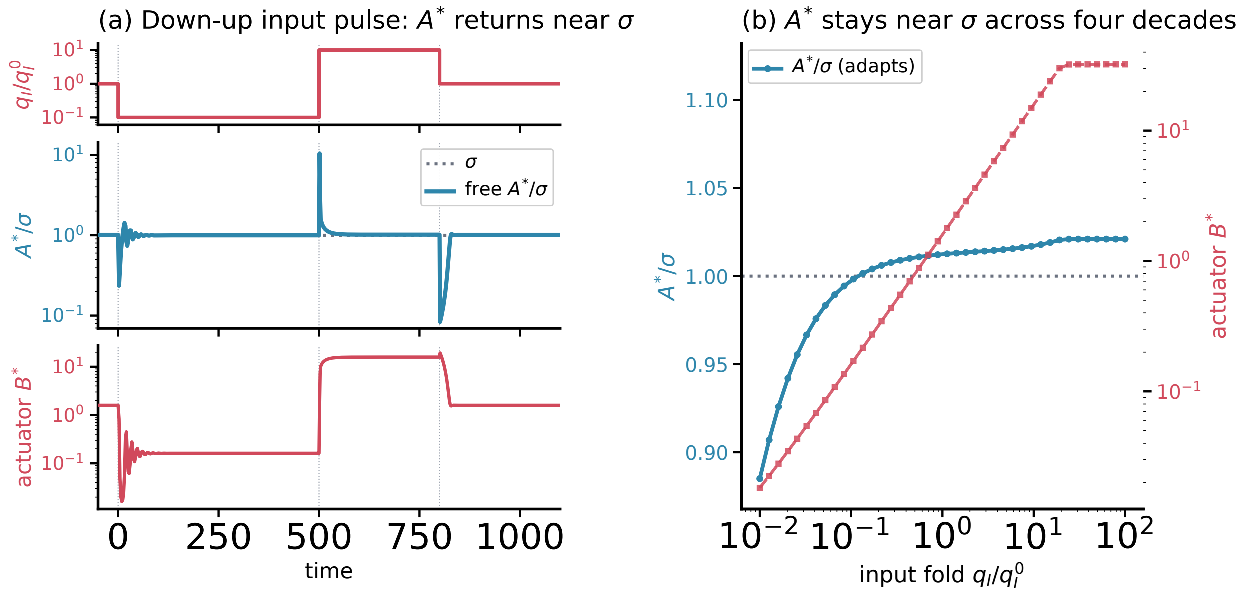

**Figure S8. A fixed-$\sigma$ JSA-box hit passes our adaptation filter.** The first parameter set found in a deterministic JSA-box search that both passes our free-$A^*$ adaptation filter and traces the literal setpoint $\sigma = k_{B2}q_E/k_{B1}$ appears only after about $2 \times 10^6$ draws under a $5 \times 10^6$-draw cap. (a) Under an input protocol $q_I/q_I^0 : 1 \to 0.1 \to 10 \to 1$, free $A^*$ returns close to the tested setpoint $\sigma$ after both downward and upward steps while the actuator $B^*$ responds strongly; $A^*$ and $B^*$ are plotted on separate axes to show the setpoint return and actuator response independently. (b) Steady-state $A^*/\sigma$ stays $\approx 1$ across four decades of input while $B^*$ tracks the input.

**Why fixed-$\sigma$ is a thin target.** The fixed-$\sigma$ condition can be read directly in flux–species variables. At any sampled steady state,

$$v_B = k_{B1}C_3 = k_{B2}C_4, \qquad k_{B1} = \frac{v_B}{C_3}, \qquad k_{B2} = \frac{v_B}{C_4}, \qquad q_E = E + C_4.$$

Therefore

$$\sigma = \frac{k_{B2}q_E}{k_{B1}} = \frac{C_3(C_4 + E)}{C_4}, \qquad A^* = \sigma \quad \Longleftrightarrow \quad A^*C_4 = C_3(C_4 + E).$$

This is an equality condition, not a dominance condition. If $C_4 \gg E$, it requires $A^* \approx C_3$. If $C_4 \ll E$, it requires $A^*C_4 \approx C_3E$. In either case, fixed-$\sigma$ adaptation is a thin slice through the broader adaptation region.

The contrast with a dominance-route setpoint is explicit. For the simple $C_4$-dominated route in (S10), use

$$K_{B1} = \frac{A^*B}{C_3}, \qquad w_B = B + B^* + C_2 + C_3 + C_4.$$

Then the post-hoc condition $A^* = \sigma K_{B1}/w_B$ is equivalent to

$$1 = \frac{C_4 + E}{C_4} \frac{B}{B + B^* + C_2 + C_3 + C_4}.$$

This condition is satisfied by the open dominance requirements $C_4 \gg E$ and free $B$ dominating the other terms in $w_B$. Thus FSS has no reason to turn fixed-$\sigma$ into a large hit class: the biologically meaningful adaptation regimes are open in species space, whereas the literal Michaelian setpoint is an additional equality slice.

The same distinction appears directly in the setpoint errors (Fig. S9). Both panels use the same all-hit populations: every free-$A^*$ adaptation hit from the JSA box and every free-$A^*$ adaptation hit from FSS. The fixed-$\sigma$ target is a poor description of the recovered hits: only 1/1027 FSS hits fall within 1% of $\sigma$. Panel (b) then asks a parallel all-hit question by comparing the same steady states with the total-pool expression $\sigma K_{B1}/w_B$. No dominance-route filter is applied before plotting this panel. This benchmark is not universal, but it captures a visible shifted setpoint class: 184/1027 FSS hits and 60/113 JSA-box hits lie within 1% of the value. Together with the explicit fixed-$\sigma$ JSA-box hit in Fig. S8, this comparison separates two possible artifacts: FSS finds genuine responsive adaptation, but it does not inflate fixed-$\sigma$ equality into a large artificial hit class.

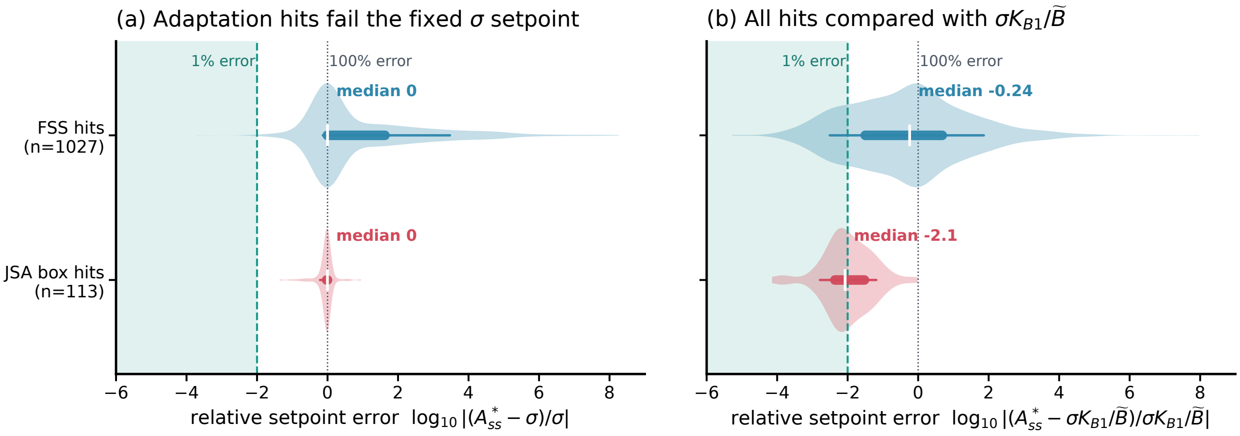

**Figure S9. Fixed-$\sigma$ is the wrong post-hoc setpoint for the recovered adaptation hits.** (a) Relative error of free $A^*$ from the literal Michaelian target $\sigma = k_{B2}q_E/k_{B1}$. The adaptation hits are input-invariant but mostly not fixed-$\sigma$ hits. (b) The same all-hit populations compared with $\sigma K_{B1}/w_B$, using the total $B$-pool of each hit. This panel is not a dominance-route subset; it shows that a shifted total-pool setpoint captures a visible class of adapting hits without making fixed-$\sigma$ adaptation generic.

#### S9 The MultiFate case study

This section assembles the MultiFate case study: the symmetry reduction that makes homogeneous-endpoint sampling two-dimensional, the protocol, and the high-dimensional asymmetric-scaling test.

The case study is organized in the same order as the adaptation case: the model and the Zhu et al. question, the sampling and filtering protocol, the all-state recovery result, a regime analysis of one recovered FSS hit, and finally the asymmetric stress test.

#### S9.1 Model setup and the all-state question

The symmetric MultiFate model uses $N$ transcription factors that share a dimerisation domain. Free monomers $A_i$ form homodimers $A_{ii} = A_i^2/K_d$, which activate the production of their own factor through a Hill term, and heterodimers $A_{ij} = 2A_iA_j/K_d$, which sequester monomer mass across factors. For a fixed total monomer pool, each scalar gene equation can have a low stable branch and a high stable branch; these branches are the OFF and ON states used in the main text. By permutation symmetry, a binary state is determined by the number $M$ of ON genes, although it represents $\binom{N}{M}$ labelled states. The case-study question is therefore whether one parameter pair $(a, b)$ can make every labelled binary pattern locally stable, i.e. all $2^N$ states.

#### S9.2 Sampling method and filtering standard

We compared three strategies. *(P1) Zhu nominal*: fix $(a, b, K_d, n) = (0.8, 20, 1, 1.5)$ and enumerate all $2^N$ binary patterns; for each, Newton-solve (10) and test linear stability. *(P2) Parameter-space sampling*: draw $(a, b)$ log-uniformly from the Zhu box ($\log_{10} a \in [-1, 0.5]$, $\log_{10} b \in [0.5, 2]$); this box contains $(0.4, 10), (0.8, 20), (1.2, 30)$; 2000 draws per $N$. *(P3) FSS sampling*: draw ON/OFF free monomer concentrations $[A_i]_{\text{high}}$ and $[A_i]_{\text{low}}$ log-uniformly in $[10^0, 10^{1.5}] \times [10^{-3}, 10^0]$. The all-OFF endpoint uses $S_{\text{off}} = N[A_i]_{\text{low}}$ and the all-ON endpoint uses $S_{\text{on}} = N[A_i]_{\text{high}}$; from those two homogeneous endpoint totals we solve the two fixed-point equations for $(a, b)$; 2000 raw endpoint draws per $N$. Endpoint pairs whose inverse map gives nonpositive $a$ or $b$ are retained as invalid draws and counted as misses. Every valid projected candidate is then judged by the same verifier as parameter sampling: one representative state for each $M = 0, \ldots, N$, Newton solve, ON/OFF classification, Jacobian stability test, and multiplication by $\binom{N}{M}$.

**Why symmetric MultiFate FSS sampling is two-dimensional.** The reduction to two sampled free monomer concentrations is a consequence of symmetry and bistability. Let $[A_i]$ be the free monomer concentration of TF $i$, let $[A_{ii}] = [A_i]^2/K_d$ be the homodimer concentration, and let $S = \sum_j [A_j]$. At steady state,

$$a + b\frac{[A_{ii}]^n}{1 + [A_{ii}]^n} = [A_i^{\text{tot}}], \qquad [A_i^{\text{tot}}] = [A_i]\left(1 + \frac{2S}{K_d}\right), \qquad [A_{ii}] = \frac{[A_i]^2}{K_d}.$$

Eliminating $[A_i^{\text{tot}}]$ gives, for fixed $S$, a scalar equation in the free monomer:

$$a + b\frac{([A_i]^2/K_d)^n}{1 + ([A_i]^2/K_d)^n} = [A_i]\left(1 + \frac{2S}{K_d}\right).$$

In the MultiFate regime this scalar equation has the standard bistable shape: a low stable root, a middle unstable root, and a high stable root. We denote the two stable free monomer roots by $[A_i]_{\text{low}}$ and $[A_i]_{\text{high}}$. Therefore, in a type-$M$ state, where $M$ genes are ON and $N - M$ genes are OFF,

$$S_M = M[A_i]_{\text{high}} + (N - M)[A_i]_{\text{low}}.$$

The manuscript projection anchors the two homogeneous endpoints. This choice is not arbitrary: any parameter set that realizes all $2^N$ labelled patterns must at least realize the all-OFF state ($M = 0$) and the all-ON state ($M = N$). The endpoint projection therefore fits the two necessary symmetric extremes first and only then uses the common verifier to test whether every mixed sector $M = 1, \ldots, N - 1$ is also stable. Their total concentrations and homodimer signals are

$$[A_i^{\text{tot}}]_{\text{high}} = [A_i]_{\text{high}}\left(1 + \frac{2N[A_i]_{\text{high}}}{K_d}\right), \qquad [A_i^{\text{tot}}]_{\text{low}} = [A_i]_{\text{low}}\left(1 + \frac{2N[A_i]_{\text{low}}}{K_d}\right).$$

With $n = 1.5$ and $K_d = 1$, define

$$h_{\text{high}} = \frac{([A_i]^2_{\text{high}}/K_d)^{1.5}}{1 + ([A_i]^2_{\text{high}}/K_d)^{1.5}}, \qquad h_{\text{low}} = \frac{([A_i]^2_{\text{low}}/K_d)^{1.5}}{1 + ([A_i]^2_{\text{low}}/K_d)^{1.5}}.$$

For a sampled pair $([A_i]_{\text{high}}, [A_i]_{\text{low}})$, the two fixed-point equations are

$$[A_i^{\text{tot}}]_{\text{high}} = a + bh_{\text{high}}, \qquad [A_i^{\text{tot}}]_{\text{low}} = a + bh_{\text{low}}.$$

Equivalently,

$$\begin{pmatrix} 1 & h_{\text{high}} \\ 1 & h_{\text{low}} \end{pmatrix} \begin{pmatrix} a \\ b \end{pmatrix} = \begin{pmatrix} [A_i^{\text{tot}}]_{\text{high}} \\ [A_i^{\text{tot}}]_{\text{low}} \end{pmatrix}.$$

Solving gives

$$a = \frac{h_{\text{low}}[A_i^{\text{tot}}]_{\text{high}} - h_{\text{high}}[A_i^{\text{tot}}]_{\text{low}}}{h_{\text{low}} - h_{\text{high}}}, \qquad b = \frac{[A_i^{\text{tot}}]_{\text{low}} - [A_i^{\text{tot}}]_{\text{high}}}{h_{\text{low}} - h_{\text{high}}}.$$

Thus homogeneous-endpoint free-monomer sampling creates many projected candidates

$$([A_i]_{\text{high}}, [A_i]_{\text{low}}) \mapsto (a, b).$$

If the inferred $a$ or $b$ is nonpositive, the raw endpoint draw has no positive parameter-space coordinate; it is kept as an invalid sample and scored as a miss. After this projection, deciding which valid $(a, b)$ values give how many stable fates is a data-processing step, not another sampling assumption: each valid candidate is verified by the same Newton solve and local-stability test as parameter-space sampling. Because the equations are permutation-symmetric, the verifier tests one representative state for each ON count $M$ and multiplies accepted representatives by $\binom{N}{M}$ labelled patterns.

##### S9.3 All-state recovery and peter-out

This subsection records the numerical counts behind Fig. 4, using scientific notation for large state spaces and reporting missed states when a draw is close to all-state recovery. Zhu et al.'s nominal pair $(a, b) = (0.8, 20)$ gives 3/4 stable states at $N = 2$: the all-OFF fixed point exists but is unstable under the deterministic local-stability criterion used here. At $N = 32$, the same nominal pair gives $529/(2^{32}) = 529/(4.29 \times 10^9)$ stable labelled states, or $1.23 \times 10^{-5}\%$ of the maximum.

The matched Zhu-box parameter search uses 2000 log-uniform draws per $N$. At $N = 32$, 0/2000 parameter draws recover all $2^{32} = 4.29 \times 10^9$ states. The best draw still misses $1.08 \times 10^8$ states, while the median draw recovers only 529 states, matching the nominal peter-out scale. The high Fig. S23 pair $(a, b) = (1.2, 30)$ is close at small $N$, missing one state at $N = 5$ and one state at $N = 6$, but the missed state is the all-OFF state and therefore the pair is not an all-$2^N$ hit.

For homogeneous-endpoint FSS at $N = 32$, 1141/2000 raw endpoint draws yield positive $(a, b)$. Counting invalid projections as misses, $899/2000 = 44.95\%$ of all raw endpoint draws realise all $2^{32} = 4.29 \times 10^9$ states. Conditional on a positive projection, the all-state rate is $899/1141 = 78.8\%$. The median raw endpoint draw misses one state. These counts support the main-text statement that FSS robustly finds the all-state regime while the fixed Zhu parameter box has zero observed all-state hits at $N = 32$.

#### S9.4 Regime analysis of a representative FSS hit

To inspect what an all-state FSS hit looks like mechanistically, we selected a representative lower-left point from the $N = 10$ all-state hits in the sampled free-monomer chart (Fig. 4f): cache column 1644, with $[A_i]_{\text{high}} = 1.270$, $[A_i]_{\text{low}} = 1.181 \times 10^{-3}$, $a = 1.209 \times 10^{-3}$, and $b = 49.893$. For a type-$M$ state,

$$S_M = M[A_i]_{\text{high}} + (N - M)[A_i]_{\text{low}}.$$

For an individual factor $i$, the total decomposes as

$$[A_i^{\text{tot}}] = [A_i] + 2[A_{ii}] + \sum_{j \neq i} [A_{ij}], \qquad [A_{ii}] = \frac{[A_i]^2}{K_d}, \qquad [A_{ij}] = \frac{2[A_i][A_j]}{K_d}.$$

The production flux decomposes as

$$v_i^+ = a + b\,h([A_{ii}]), \qquad h([A_{ii}]) = \frac{[A_{ii}]^n}{1 + [A_{ii}]^n}.$$

Figure S10 plots the corresponding fractions for $M = 0, \ldots, 10$. ON genes are sustained by the Hill production term and become increasingly heterodimer-buffered as more genes are ON. OFF genes keep nearly all positive production in the basal term, while their total is rapidly absorbed into heterodimers once any ON factor is present. The flux dominance is separate from the species composition dominance. For this selected hit,

$$\frac{bh(A_{ii}^{\text{on}})}{a + bh(A_{ii}^{\text{on}})} = 0.999964, \qquad \frac{a}{a + bh(A_{ii}^{\text{off}})} = 0.999932.$$

Thus ON states are high because the inducible Hill flux dominates, whereas OFF states stay low because basal production dominates even when the OFF monomer pool is mostly sequestered into heterodimers by nearby ON factors. This is the dominance-regime reading of one concrete FSS hit: all-state recovery is not a single numerical accident, but a structured balance between ON autoregulatory production, OFF basal production, and heterodimer sequestration.

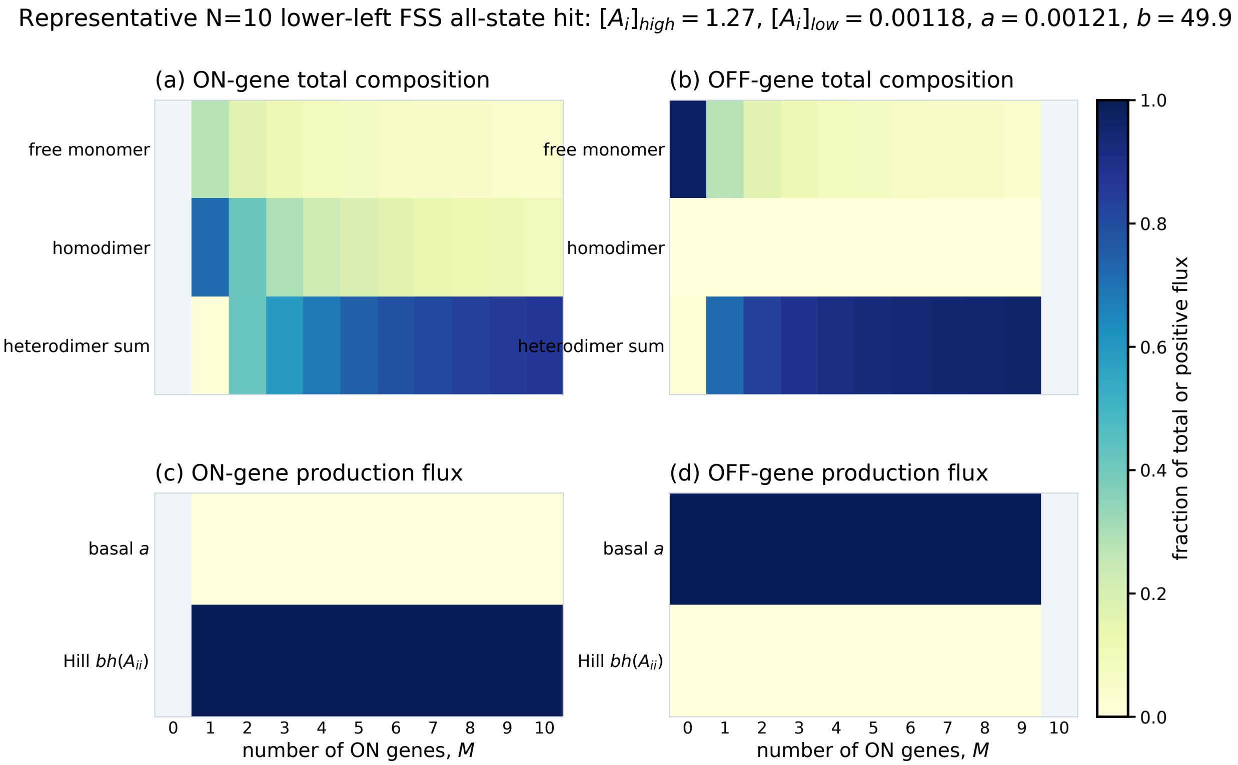

**Figure S10. Species and flux dominance in a representative $N = 10$ all-state FSS hit.** The selected point is a lower-left all-state hit in the free-monomer endpoint chart of Fig. 4f. For each ON count $M$, panels (a,b) show the fraction of an ON or OFF factor total carried by free monomer, homodimer, or the sum of heterodimers; panels (c,d) show the fraction of positive production flux carried by basal production $a$ or the Hill term $bh(A_{ii})$. In this hit, the ON flux is essentially Hill-dominated and the OFF flux is essentially basal-dominated, while the species totals shift with $M$ through heterodimer sequestration. Blank columns mark absent classes: there are no ON genes at $M = 0$ and no OFF genes at $M = N$.

#### S9.5 High-dimensional scaling: asymmetric MultiFate

The symmetric MultiFate result could, in principle, depend on the two-dimensional collapse created by permutation symmetry. We therefore also run a deliberately asymmetric stress test in which each transcription factor $i$ has its own production pair $(a_i, b_i)$, while all factors still share the same dimerisation constants $(K_d, n)$. A parameter-space candidate is then the $2N$-vector $(a_1, \ldots, a_N, b_1, \ldots, b_N)$, sampled directly from the per-factor Zhu box. The endpoint FSS candidate is the $2N$-vector of species endpoints $(\ell_1, \ldots, \ell_N, h_1, \ldots, h_N)$, where $\ell_i$ is the intended OFF free-monomer level and $h_i$ is the intended ON free-monomer level.

The inverse map is still local in genes after the two endpoint free-monomer sums are fixed:

$$S_{\text{off}} = \sum_i \ell_i, \qquad S_{\text{on}} = \sum_i h_i.$$

For each $i$, the OFF and ON fixed-point equations give a $2 \times 2$ linear solve for $(a_i, b_i)$. Endpoint draws that return any nonpositive $(a_i, b_i)$ are rejected by this positivity filter; accepted candidates are then evaluated by the same $N$-dimensional Newton solve and Jacobian stability test used for parameter-space sampling.

Because asymmetric shared sequestration makes literal all-$2^N$ co-stability much harder than in the symmetric case, the diagnostic in Fig. S11 is the number of labelled ON/OFF patterns realised by each accepted candidate. This metric separates two effects: the raw endpoint window has a positivity-filter cost, but conditional on passing that algebraic filter, endpoint FSS lands in high-co-stability regions far more often than direct parameter sampling. At $N = 15$, only 1.7% of raw endpoint draws are accepted, but 183/300 accepted FSS candidates realise all 32768 labelled states and the mean accepted count is 32767.4; parameter-space sampling has mean count 53.8 under the same verifier.

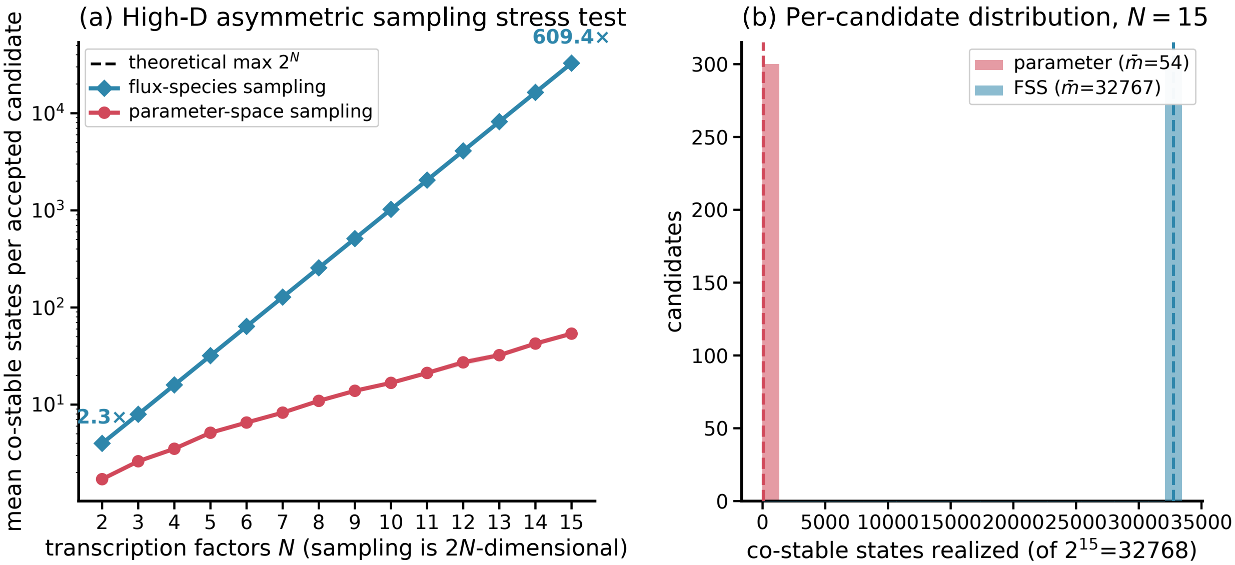

**Figure S11. Asymmetric MultiFate stress test.** The symmetry reduction is removed by giving each gene its own $(a_i, b_i)$. Parameter sampling draws these $2N$ parameters directly from the per-factor Zhu box. Endpoint FSS draws OFF/ON free-monomer endpoints $(\ell_i, h_i)$, solves for $(a_i, b_i)$, keeps only positive projections, and then applies the same $N$-dimensional stability verifier. (a) Mean number of co-stable labelled ON/OFF patterns among accepted candidates. (b) Distribution of co-stable-state counts at $N = 15$, showing the conditional advantage of the endpoint chart after the positivity filter.
